# Rapid associative spine enlargement is required for cognitive function and stable wakefulness

**DOI:** 10.1101/2025.11.06.686434

**Authors:** Siqi Zhou, Takeshi Sawada, Hitoshi Okazaki, Tomoki Arima, Shunki Takaramoto, Sadam Khan Panezai, Masanari Ohtsuka, Shin-Ichiro Terada, Masashi Kondo, Takaaki Hashimoto, Kenichi Ohki, Masanori Matsuzaki, Sho Yagishita, Haruo Kasai

## Abstract

Awake cognition is usually attributed to arousal systems and ongoing electrical activity, yet these do not explain how cortical processing is coupled to prior context and memory formation, Associative synaptic plasticity provides a cellular mechanism for such coupling, but is viewed as memory-related rather than an online requirement for cognition. Here we developed SynC, an A/C-activated chemogenetic perturbation that selectively and reversibly blocks associative spine enlargement while sparing non-associative spine dynamics, excitability, synaptic transmission and NMDA receptor-mediated responses. In mice with broad frontoparietal SynC expression, acute A/C reversibly impaired open-field centre exploration, abolished laser-dot chasing, and delayed feeding initiation. During quiet-wake immobile epochs, cortical firing rates and γ-band power were preserved, but pairwise spike correlations were reduced. Mice also intermittently showed sudden behavioural arrest with reduced γ activity but without the robust δ elevation of slow-wave sleep. Critically, in vivo two-photon glutamate uncaging induced seconds-scale associative spine enlargement in ∼24% of neocortical spines, acutely blocked by SynC–A/C. These A/C-induced behavioural, circuit and spine-enlargement effects recovered within 1 h. Thus, wake-like firing and γ activity are not sufficient to sustain awake cognitive function. Rapid associative spine enlargement is required to maintain functional cortical coupling and stable wakefulness.

## Introduction

Wakefulness is defined operationally by EEG and EMG criteria, and awake cognition is usually attributed to arousal systems and the ongoing electrical activity they sustain. Yet moment-to-moment cortical processing is not merely a sequence of responses to current input; it depends on relating ongoing activity to prior context and, in many cases, to memory formation^1–3^. Associative synaptic plasticity provides a cellular mechanism for such context-dependent change, but it has generally been viewed as a mechanism for learning and memory rather than as an online requirement for cognition.

Pharmacological studies have implicated NMDA receptor–dependent processes in learning and cognition; however, NMDA receptor antagonists broadly suppress NMDA receptor–mediated synaptic currents and alter baseline neurophysiology, complicating causal attribution to rapid associative plasticity per se^4,5^. Thus, whether rapid associative plasticity, distinct from general synaptic transmission, is required for cognitive function and stable wakefulness remains unresolved.

A central experimental obstacle has been to isolate rapid associative synaptic plasticity from general synaptic transmission and baseline neuronal activity. Most excitatory synapses reside on actin-rich dendritic spines, and coincident pre- and postsynaptic activity can induce a rapid enlargement of spine heads that persists beyond the initial induction period^6–8^. We refer to this coincidence-dependent structural plasticity as associative spine enlargement. This process is closely associated with synaptic potentiation and synaptic coupling^9^. Its onset, however, has not been resolved on the timescale of cognition: induction and imaging protocols have typically spanned tens of seconds to minutes even in slice preparations, leaving untested whether associative spine enlargement can act within ongoing cortical processing. Directly testing whether it is required online in awake cortex has been difficult because rapid associative plasticity must be separated from short-term presynaptic plasticity, ongoing baseline fluctuations^10^, and whole-cell recording washes out the plasticity itself^11,12^.

Testing moment-to-moment necessity requires acute, reversible loss of function; gain-of-function or chronic manipulations alone are insufficient. We therefore built on the synaptic chemogenetic strategy established with SYNCit-K (SynK), which promotes spine enlargement^13^, to generate SYNCit-C (SynC), an A/C-activated postsynaptic perturbation designed to block activity-dependent dendritic spine enlargement rather than general excitatory transmission. SynC acutely and reversibly blocks spine enlargement while sparing basal synaptic transmission and intrinsic membrane properties, minimizing confounds from cortical silencing. This capability distinguishes SynC from classical manipulations of NMDA receptor signalling and downstream kinases: CaMKII perturbations impair persistent components of structural potentiation but do not abolish the fast transient spine enlargement^14,15^, whereas SynC blocks the enlargement itself while preserving NMDA receptor–mediated responses. Using SynC, together with an in vivo neocortical uncaging assay developed here, we asked whether rapid associative spine enlargement is required online during wakefulness.

### Selective and reversible inhibition of spine enlargement by SynC

We designed SynC to block activity-dependent spine enlargement by exploiting the ability of a rapamycin derivative (A/C) to irreversibly induce heterodimerization between FKBP (FK506-binding protein) and FRB (the FKBP12–rapamycin-binding domain of mTOR) (Fig. 1a)^13,16^. Because SynK implicated Rac1 activation by Rac1-GEFs in spine enlargement (Fig. 1b)^17^, we generated SynC in which both components were expressed from the CaMKII promoter: a PSD-anchored PSDΔ1,2-FRB and a cytosolic FKBP-Rac1-GAP (the GAP domain of α1-chimaerin) to inactivate Rac1 by GTP hydrolysis (Fig. 1c). PSDΔ1,2-FRB-mVenus localized selectively to postsynaptic densities and was absent from axons and somas (Extended Data Fig. 1a,b).

**Fig. 1.**
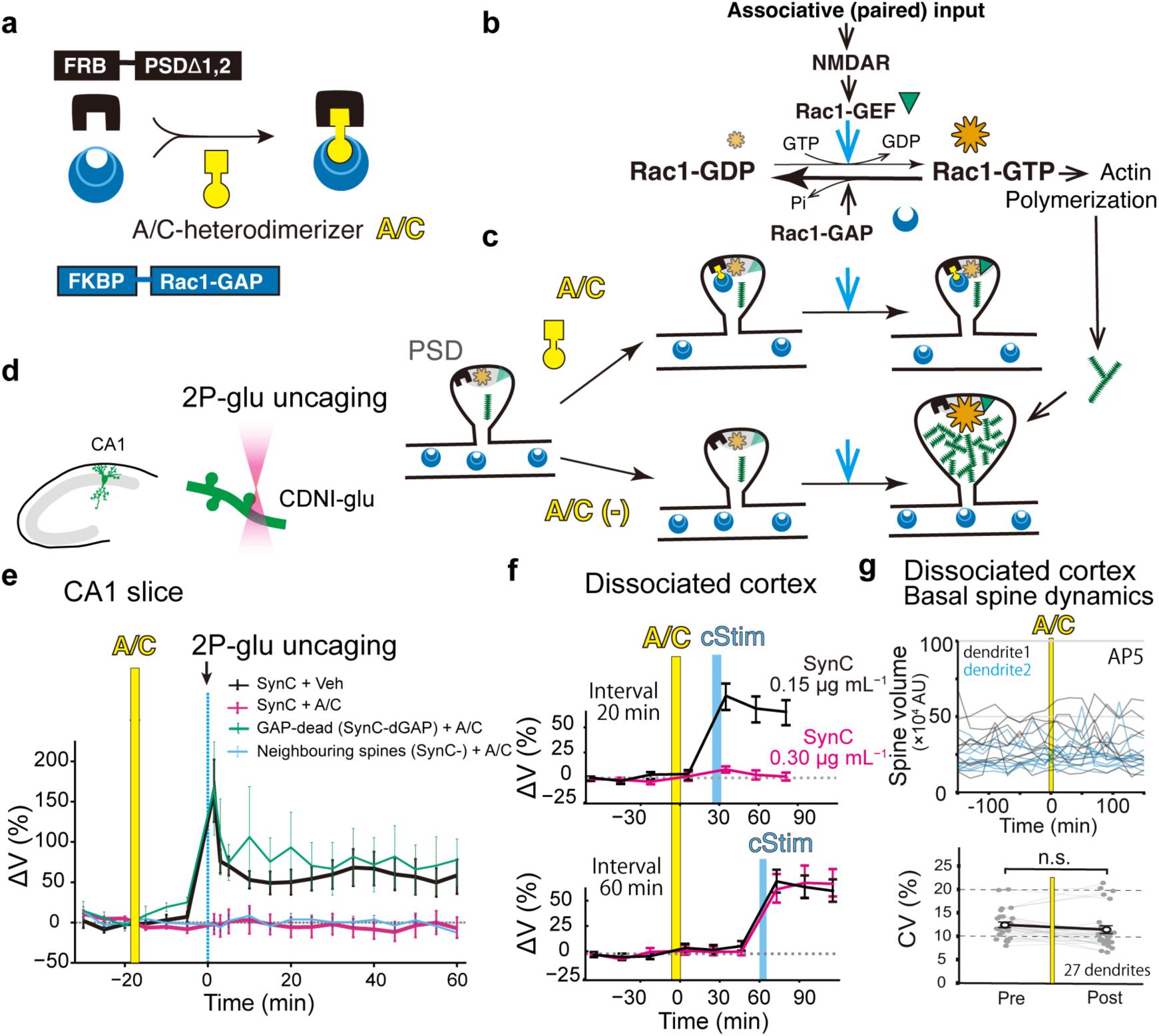
A/C-triggered SynC reversibly and selectively blocks associative spine enlargement. **a**, The A/C heterodimerizer induces FRB–FKBP dimerization. **b**, Associative input activates NMDAR and Rac1-GEF, which loads Rac1 with GTP, causing actin polymerization, whereas a GAP quickly inactivates Rac1 by catalysing GTP hydrolysis. **c**, SynC design. FRB is anchored at the postsynaptic density (PSD) via PSDΔ1,2, and FKBP-Rac1-GAP is cytosolic. Without A/C, endogenous Rac1 signalling permits enlargement; A/C recruits Rac1-GAP to the PSD, suppressing Rac1-GTP and preventing enlargement. **d**, Two-photon uncaging of CDNI-glutamate at a single spine in organotypic rat hippocampal slice cultures (CA1) transduced with AAV::SynC (Extended Data Fig. 2a). **e**, In SynC-expressing spines, uncaging-evoked spine enlargement is blocked by A/C but not by vehicle. Enlargement is unaffected in the GAP-dead control (SynC-dGAP) and in neighbouring SynC-negative spines exposed to A/C. Exact statistics are provided in Extended Data Fig. 2b. **f**, SynC completely blocks spine enlargement induced by chemical stimulation (cStim) in culture at 0.30 µg mL^−1^ (DNA concentration, magenta) but not at 0.15 µg mL^−1^ (black). Top, blockade tested 20 min after A/C. Bottom, the blockade had recovered by ∼60 min after A/C (see Extended Data Fig. 3). Exact statistics are provided in Extended Data Fig. 3a–c. **g**, Basal (non-associative) spine dynamics in the presence of AP5 (50 µM) are spared by SynC activation. Top, representative traces of spine-volume fluctuations (AU) in SynC-expressing spines, aligned to A/C application; black and blue traces are from two different dendrites. Bottom, coefficient of variation (CV) of spine volume before and after A/C in the same dendrites (27 dendrites, 268 spines, 10 cells, 4 cultures). Paired *t*-test, *t*(26) = −1.57, *P* = 0.13.

Application of A/C rapidly recruited the FKBP fusion from dendritic shafts into spines (Extended Data Fig. 1c–f and 2a) and abolished rapid spine enlargement evoked either by two-photon glutamate uncaging in 0 Mg^2+^ ACSF (Fig. 1e; statistics in Extended Data Fig. 2b)^6,18^ or by a chemical stimulation protocol (cStim, Fig. 1f and Extended Data Fig. 1c–f), an effect not observed with the GAP-dead mutant SynC-dGAP (Fig. 1e and Extended Data Fig. 1c–f). These results indicate that spine-localized Rac1-GAP catalysis is sufficient to suppress Rac1-dependent spine enlargement under these conditions^19^.

Basal excitatory and inhibitory synaptic transmission and intrinsic membrane properties were unchanged, and NMDA receptor–mediated slow EPSCs were unaffected (Extended Data Fig. 2c–g), consistent with stable somatic and spine Ca^2+^ signals in vivo (see Extended Data Fig. 9d, e). Postsynaptic SynC expression also spared non-associative short-term presynaptic plasticity^20,21^, including facilitation, depression and post-tetanic potentiation (Extended Data Fig. 2h,i).

For most subsequent experiments, we used a bicistronic plasmid in which the FRB and FKBP modules of SynC were co-expressed via an internal ribosome entry site (IRES), enabling single-vector AAV-mediated delivery for in vivo use (Extended Data Fig. 3a)^13^. The A/C-induced suppression of spine enlargement was transient, dissipating within ∼1 h at the low expression levels used in subsequent experiments (Fig. 1f; Extended Data Fig. 3a–c). A/C action outlasted 1 h at higher SynC expression (Extended Data Fig. 3c), indicating that the recovery time reflects expression level. Functional recovery despite persistent spine enrichment of FKBP-mVenus suggests functional attenuation of the artificially PSD-tethered Rac1-GAP module rather than loss of spine recruitment. This transient action provided an internal recovery control across subsequent cellular, circuit and behavioural experiments. SynC did not alter NMDA-evoked spine shrinkage (Extended Data Fig. 3d), nor baseline spine morphology or intrinsic spine-volume fluctuations, which include spontaneous enlargements and reflect ongoing actin-based spine motility^22^ (Fig. 1g; Extended Data Fig. 3e), indicating that SynC selectively blocks associative spine enlargement. For in vivo AAV delivery, mVenus was replaced by an HA tag, reducing the expression cassette to 4.5 kb (Fig. 2a, bottom).

**Fig. 2.**
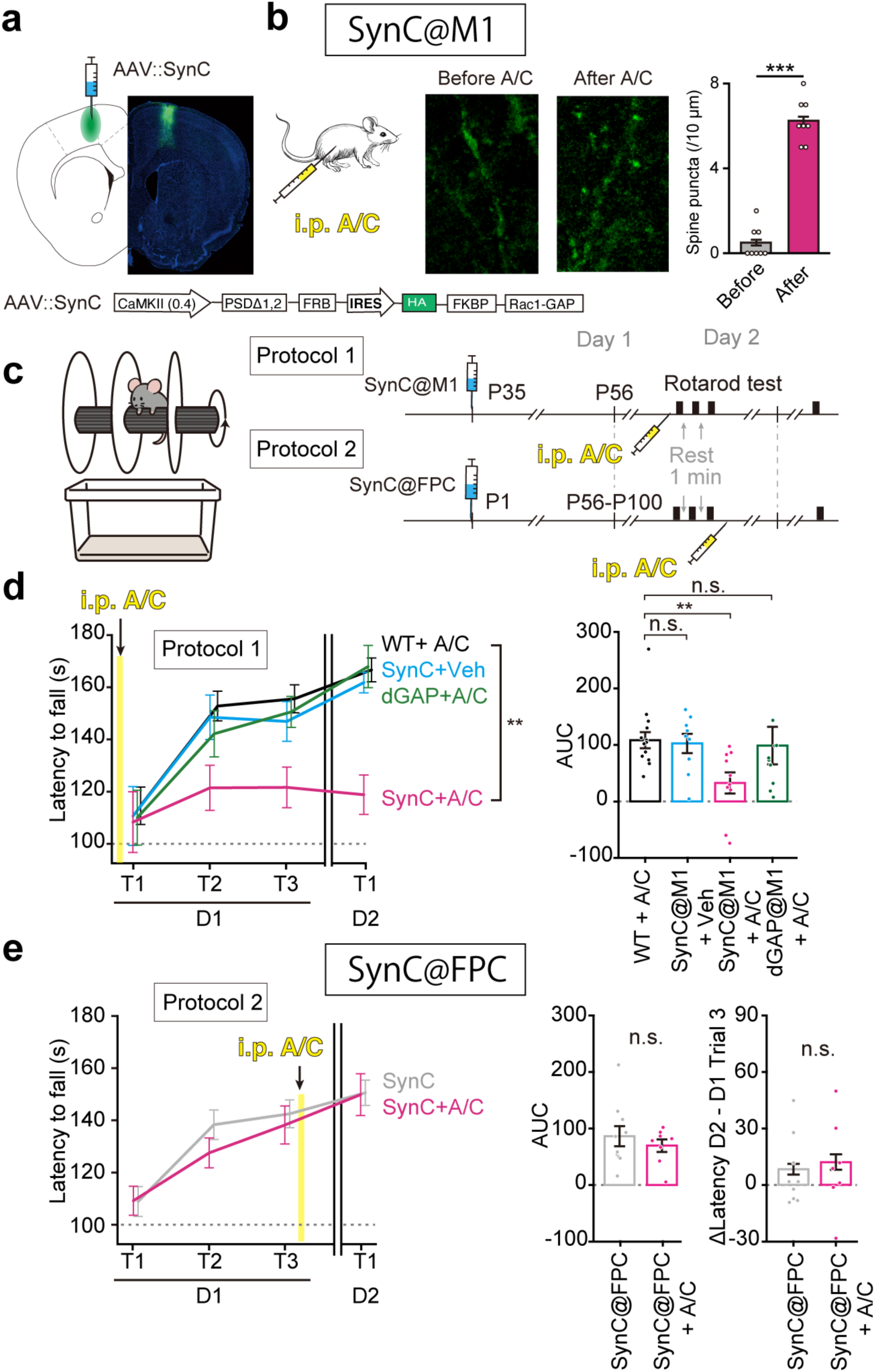
Local SynC in M1 blocks motor learning without affecting initial performance or consolidation. All behavioural tests in Fig. 2 were performed during the light (inactive) phase. **a**, Bilateral injection of AAV::SynC into M1 and representative coronal section showing expression. Bottom, bicistronic SynC construct with HA tag. **b**, Immunohistochemical detection of the HA tag in M1 dendrites from separate mice, untreated or 30 min after i.p. A/C. Right, HA-positive spine puncta per 10 µm (Methods, 8 dendritic segments from 4 mice [before] and 8 dendritic segments from 2 mice [after]). Mann–Whitney, *U* = 0, *P* = 1.6 × 10^−4^. **c**, Rotarod task and experimental protocols. Protocol 1, SynC injected into M1 in adulthood; A/C before training (acquisition test). Protocol 2, SynC injected into FPC in pups; A/C after Day 1 training (post-training/performance test). **d**, Protocol 1 in SynC@M1. Left, learning curves showing latency to fall; right, AUC. Group sizes: *n* = 15 WT + A/C, 8 SynC@M1 + vehicle, 10 SynC@M1 + A/C, and 11 SynC-dGAP@M1 + A/C mice. Kruskal–Wallis, *H*(3) = 9.26, *P* = 0.026; Steel’s post hoc vs WT + A/C: SynC@M1 + vehicle, *P* = 0.85; SynC@M1 + A/C, *P* = 0.0082; SynC-dGAP@M1 + A/C, *P* = 0.62. Initial performance (first trial after A/C): Kruskal– Wallis, *H*(3) = 0.65, *P* = 0.88. **e**, Protocol 2 in SynC@FPC: overnight improvement was unaffected when A/C was administered after the initial training session (*n* = 10 A/C[−], 8 A/C[+]). Error bars, s.e.m. Mann–Whitney (A/C after Day 1 training): AUC, *U* = 44, *P* = 0.76; Δlatency, *U* = 32.5, *P* = 0.53. Error bars, s.e.m.

### Local SynC preserves visual responses and blocks motor learning

We tested whether SynC perturbs basic response properties in primary visual cortex (V1). SynC was delivered to Thy1–GCaMP6s transgenic mice by AAV (1.2 × 10^13^ gc mL^−1^; 1 µL; Extended Data Fig. 4a,b). Four weeks later, we recorded grating-evoked responses from hundreds of neurons. A/C (2 mg kg^−1^) was then administered intraperitoneally (i.p.) under light anaesthesia (Methods). Retinotopy, orientation selectivity, and neuronal firing rates were unchanged after A/C in SynC-expressing mice (SynC@V1) (Extended Data Fig. 4c–k). Thus, acute SynC activation spared basic visual processing, consistent with the intact synaptic transmission and intrinsic membrane properties measured in vitro (Extended Data Fig. 2).

We next examined motor learning in primary motor cortex (M1)^23^. SynC was bilaterally delivered to the forelimb representation by AAV (Fig. 2a), and rotarod training was performed 4 weeks later. Within 30 min of i.p. A/C, translocation of HA-FKBP-Rac1-GAP to dendritic spines was detected in fixed SynC@M1 tissue (Fig. 2b). In controls, performance improved rapidly within the first 20 min, typically plateaued by ∼30 min, and was maintained on the next day (Fig. 2c and d). In SynC@M1 mice, A/C abolished rotarod learning, whereas A/C alone or A/C in SynC-dGAP mice left learning and consolidation intact (Fig. 2d). Initial performance (latency to fall at the first trial after A/C) did not differ across groups (Fig. 2d), indicating that A/C selectively abolished the progressive improvement across training sessions rather than baseline motor ability. These data indicate that motor learning in this task requires Rac1-dependent spine enlargement which is blocked by SynC–A/C at spines.

### Frontoparietal SynC impairs goal-directed behaviour

To test the impact of spine enlargement on goal-directed behaviour^24^, we bilaterally transduced SynC using AAV (1.2 × 10^13^ gc mL^−1^; 4 µL) in neonatal pups (P1) under hypothermic anaesthesia, targeting a site between S1 and M2 in the sagittal plane (AP, 0 to 1 mm; ML, 1 mm) (Fig. 3a; Extended Data Fig. 5a,b). Injection at P1 ensured extensive cortical distribution^25^. At 6 to 12 weeks post-injection, histology showed broad expression spanning from prefrontal cortex (M2) to posterior parietal cortex (PPC), hereafter frontoparietal cortex (FPC) (Extended Data Fig. 5c; Methods). Across transduced areas, 20 to 80% of NeuN^+^ layer 5 neurons expressed SynC, varying by cortical region (Extended Data Fig. 5c). No gross or cytoarchitectural abnormalities were detected (Fig. 3a), and SynC@FPC mice displayed rotarod performance comparable to WT mice (Fig. 2e). Expression was most prominent in layer V neurons, moderate in layers II/III, and nearly absent in layer VI, consistent with the CaMKII promoter (Fig. 3a; Extended Data Fig. 5d). PSDΔ1,2-FRB-mVenus was largely restricted to dendritic spines, with no labelling in dendritic shafts, somatic cytosol (Fig. 3b), nucleus, or axons in vivo (Extended Data Fig. 5e).

**Fig. 3.**
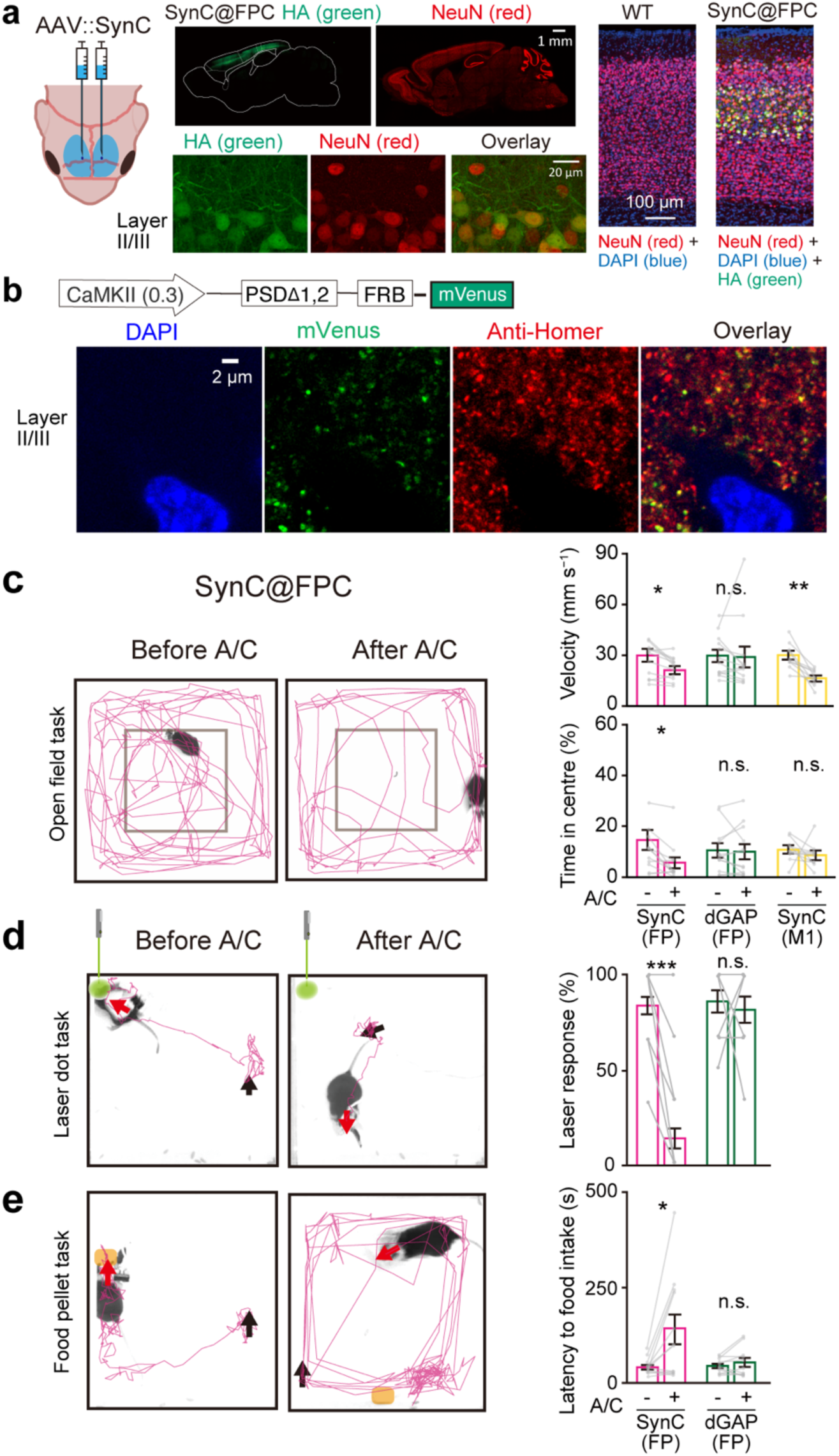
SynC activation in frontoparietal cortex disrupts goal-directed behaviour. All behavioural tests in Fig. 3 were performed during the light (inactive) phase. **a**, Neonatal bilateral cortical injection of AAV::SynC and representative sagittal sections at 10 weeks. SynC was detected by anti-HA (green) and neurons by anti-NeuN (red); DAPI (blue). Right, cortical columns from WT and SynC@FPC mice. Scale bars, 1 mm (sagittal sections), 20 µm (layer II/III) and 100 µm (right). **b**, Immunohistochemistry of an 8-week-old mouse injected with AAV::PSDΔ1,2-FRB-mVenus, showing DAPI, mVenus, Alexa-conjugated anti-Homer antibody and their overlay. SynC-FRB (construct at top) was localized only to dendritic spines and was excluded from dendritic shafts and somata. Scale bar, 2 µm. **c**, Open-field trajectories before and after i.p. A/C. Quantification of running velocity and time spent in the centre. Group sizes: *n* = 12 SynC@FPC, 12 SynC-dGAP@FPC, and 10 SynC@M1 mice. Bonferroni-corrected Wilcoxon signed-rank tests (velocity): SynC@FPC, *W* = 6, *P* = 0.0205; SynC-dGAP@FPC, *W* = 20, *P* = 0.454; SynC@M1, *W* = 0, *P* = 0.00586. (time in centre): SynC@FPC, *W* = 6, *P* = 0.0205; SynC-dGAP@FPC, *W* = 38, *P* = 1.0; SynC@M1, *W* = 10, *P* = 0.252. **d**, Laser-dot chasing task. Representative trajectories and laser response rate (*n* = 26 SynC@FPC and 11 SynC-dGAP@FPC mice). Bonferroni-corrected Wilcoxon signed-rank tests: SynC@FPC, *W* = 0, *P* = 1.79 × 10^−5^; SynC-dGAP@FPC, *W* = 6, *P* = 1.0. **e**, Food-pellet initiation task. Representative trajectories and latency to feeding initiation (*n* = 12 SynC@FPC and 11 SynC-dGAP@FPC mice). Bonferroni-corrected Wilcoxon signed-rank tests: SynC@FPC, *W* = 10, *P* = 0.042; SynC-dGAP@FPC, *W* = 23, *P* = 0.826. Time spent in State-C was excluded from the open-field velocity and feeding-latency measurements. For c–e, analyses were performed in animals meeting predefined State-C criteria. Error bars, s.e.m.

Following i.p. A/C, SynC@FPC mice showed marked impairments in goal-directed behaviour (Fig. 3c–e). In the open field, centre exploration was reduced in SynC@FPC mice but not in SynC@M1 mice, despite reductions in locomotor speed in both groups (Fig. 3c). In a laser-dot chasing assay (Methods), the laser response rate dropped from 80% pre–A/C to 15% post–A/C (*n* = 26 SynC@FPC; Fig. 3d; Extended Data Movies 1 to 4), whereas SynC-dGAP@FPC controls were unaffected (*n* = 11; Fig. 3d). In a food-pellet task, SynC–A/C increased the latency to initiate feeding from ∼30 s to ∼150 s despite frequent pellet encounters. Once the pellet was grasped, however, consummatory behaviours proceeded normally (Fig. 3e; Extended Data Movies 5 to 7). Remarkably, this selective delay in feeding initiation despite preserved consummatory behaviour closely parallels the classic appetitive–consummatory dissociation observed after chronic decerebration in rats^26^. To test whether these goal-directed deficits persisted during the animals’ active phase and to resolve their temporal profile, we examined a separate dark-phase cohort. After A/C, SynC@FPC mice continued to locomote at velocities comparable to WT mice (Extended Data Fig. 6) but failed to reduce their distance to either the laser dot or the food pellet (see Fig. 4g; Extended Data Fig. 6b,e), in contrast to the reduced open-field speed measured without an eliciting stimulus in the light phase (Fig. 3c).

**Fig. 4.**
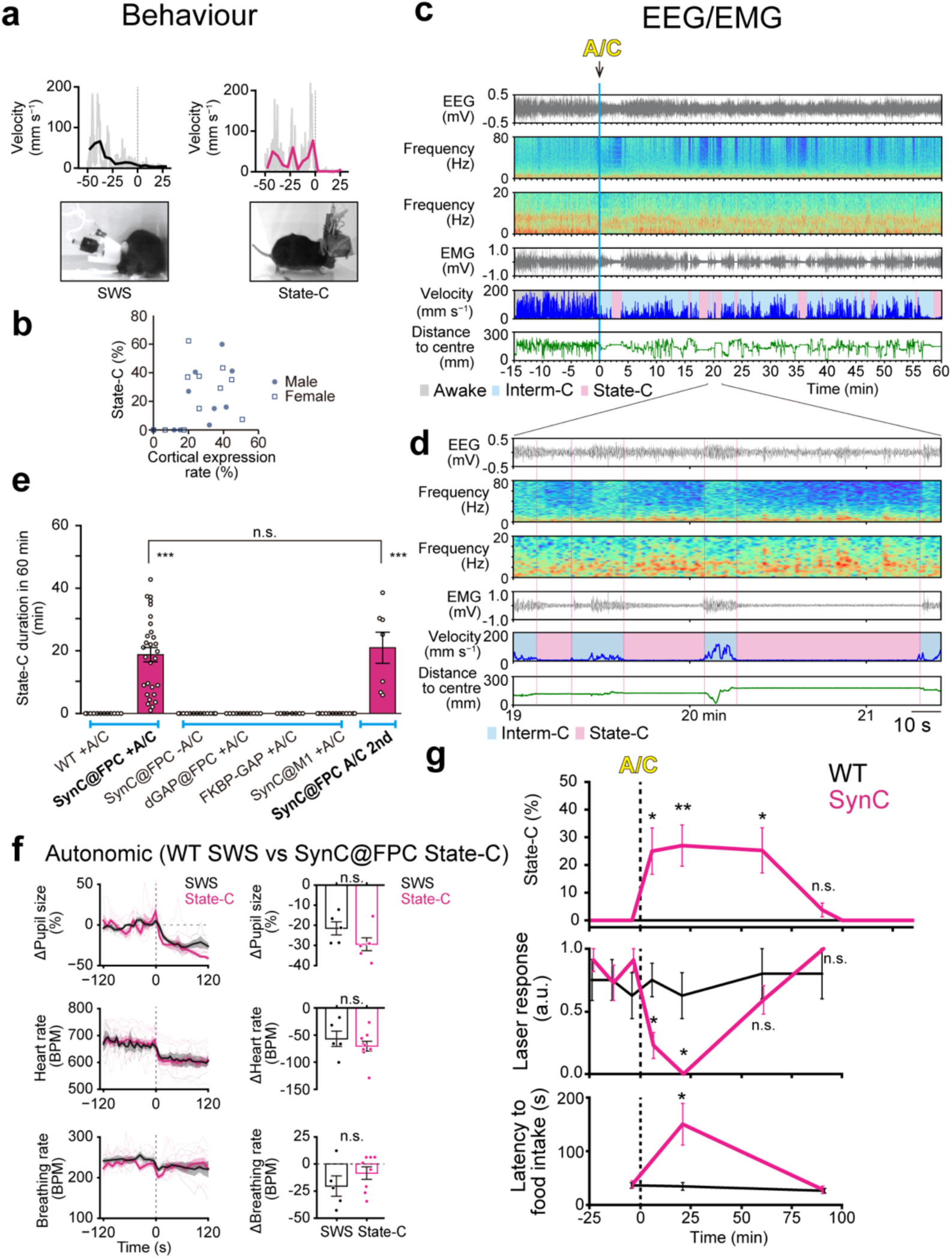
Definition and physiological signature of State-C. Recordings in Fig. 4 were performed during the dark (active) phase. **a**, Example transitions from Awake into slow-wave sleep (SWS) in wild-type (WT) mice and from Interm-C into State-C in SynC@FPC mice, showing locomotor velocity and posture. **b**, Cortical SynC expression vs State-C (%) duration within 1 h after A/C (*n* = 21 mice); mice with <20% expression show no State-C (*n* = 5). Filled and open symbols denote male and female mice, respectively. **c**, Representative EEG/EMG recordings, power spectra, velocity, distance to centre and state classification before and after i.p. A/C. **d**, Interm-C→State-C transition showing abrupt suppression of γ-band power and spindle-like events without a sustained increase in δ power. **e**, Cumulative time spent in State-C during the first hour after A/C across groups. dGAP stands for SynC-dGAP. WT + A/C (*n* = 12), SynC@FPC + A/C (*n* = 29), SynC@FPC - A/C (*n* = 29), dGAP@FPC+A/C (*n* = 11), FKBP-GAP + A/C (*n* = 8), SynC@M1+A/C (*n* = 14), SynC@FPC A/C 2nd (*n* = 7). Mann-Whitney U test: *U* = 0, *P* = 4.78 × 10^−7^, WT + A/C vs SynC@FPC + A/C; *U* = 85, *P* = 0.522, SynC@FPC + A/C vs SynC@FPC + A/C 2nd. **f**, Autonomic changes during transitions into State-C (pupil and heart rate, *n* = 12 transitions each; breathing rate, *n* = 20 transitions) compared with WT SWS transitions (*n* = 10). Mann–Whitney tests (State-C vs WT SWS transitions): pupil, *U* = 70, *P* = 0.54; heart rate, *U* = 68, *P* = 0.63; breathing rate, *U* = 74, *P* = 0.27. **g**, Time course after A/C showing State-C occupancy (*n* = 9 WT, 8 SynC mice), laser-dot responses (*n* = 5–8 WT, 8–11 SynC mice), and latency to feeding initiation (*n* = 8 WT, 7 SynC mice). State-C occupancy (Mann–Whitney vs WT with Bonferroni correction across the four time points): *U* = 9, *P* = 0.011 (0–10 min); *U* = 4.5, *P* = 0.0034 (10–30 min); *U* = 13.5, *P* = 0.034 (50–70 min); *U* = 27, *P* = 0.58 (70–90 min). Laser-dot responses (Mann–Whitney vs WT with Bonferroni correction across the four time points): *U* = 73, *P* = 0.048 (*n* = 8 WT, 11 SynC, 0–10 min); *U* = 71.5, *P* = 0.014 (*n* = 8 WT, 11 SynC, 10–30 min); *U* = 37, *P* = 1.0 (*n* = 5 WT, 11 SynC, 50–70 min); *U* = 16, *P* = 1.0 (*n* = 5 WT, 8 SynC, 70–90 min). Latency to feeding initiation (Mann– Whitney vs WT with Bonferroni correction across the two time points): *U* = 8, *P* = 0.041 (0–30 min); *U* = 32, *P* = 1.0 (60–90 min). Time spent in State-C was excluded from feeding-latency measurements. Error bars, s.e.m.

### SynC induces intermittent State-C

Following i.p. injection of A/C in the open-field arena, 41 of 57 SynC@FPC mice exhibited intermittent behavioural arrests beginning 5 to 15 min post-injection (Fig. 4a). These arrests cumulatively occupied ∼31.2% (± 3.8%) of the first hour. Arrest episodes frequently occurred during quadrupedal standing at the end of running bouts (“standing”), while navigating corners (“corner”), or during grooming (“grooming”) (Extended Data Fig. 7; Extended Data Movies 8 to 11). During these episodes, animals remained motionless in a low-arousal arrest state with eyes open, head elevated, and tail extended, generally retaining the posture in which arrest occurred, except for occasional minor postural shifts in the “corner” state (Extended Data Fig. 7b; see below). We refer to these behavioural-arrest episodes as State-C, and to the intervening post-A/C periods outside State-C before recovery as Interm-C.

To relate phenotype to SynC expression levels, we performed whole-brain histology in 21 of the 32 mice. The five mice without State-C showed <20% neuronal expression whereas State-C appeared when expression exceeded this threshold (Fig. 4b; Extended Data Fig. 5c). Accordingly, behavioural analyses in Fig. 3c–e were restricted to mice exhibiting State-C, which served as an operational indicator of sufficient cortical SynC expression. State-C required Rac1-GAP activity in neocortical dendritic spines, as A/C did not induce State-C in SynC-dGAP@FPC mice with histologically confirmed expression (Fig. 4e; Extended Data Fig. 5b, c). A/C also failed to induce State-C in mice expressing HA-FKBP-Rac1-GAP without the PSDΔ1,2-FRB anchor (Fig. 4e), making endogenous mTOR signalling an unlikely confound. Local SynC expression in M1 never caused State-C (Fig. 4e).

Electrophysiologically, State-C was characterized by reduced EMG tone and suppressed γ-band power (Fig. 4c,d; Extended Data Fig. 8a–c). State-C did not exhibit a robust increase in slow-wave (δ-band) power, distinguishing it from SWS (Fig. 4c,d; Extended Data Fig. 8c). This contrasts with the robust slow-wave increase previously reported for SynK, which promotes spine enlargement in frontal cortex^13^. Isolated sleep spindles, a hallmark of light sleep, were frequently observed in State-C but rarely seen in matched control recordings (Extended Data Fig. 8a)^27^. Spike-and-wave discharges characteristic of absence seizures^28^ were not observed (Extended Data Movie 8). Pupils constricted (miosis), heart rate modestly decreased, and respiration remained unchanged, closely matching the changes measured during WT SWS transitions (Fig. 4f). Notably, State-C was not accompanied by a sleep posture: mice retained the posture in which arrest occurred, with eyes open, head elevated and tail extended, and episodes were distributed across the arena, whereas SWS occurred in a curled posture concentrated near corners (Extended Data Fig. 7b,c). This autonomic and motor signature differed from arrest induced by fear^29^ or brainstem stimulation^30^. REM sleep (θ-band dominance with low EMG; Extended Data Fig. 8a) was not observed during State-C, distinguishing it from narcolepsy^31^. For differential classification from sleep, seizure, anaesthesia, and related arrest states, see Supplementary Table 1.

EEG γ-band power (30 to 80 Hz) in Interm-C remained comparable to awake periods recorded from the same mice prior to A/C administration (awake baseline) (Fig. 4c, d; Extended Data Fig. 8c). The duration of Interm-C bouts followed a heavy-tailed distribution, similar to awake bouts in WT mice, whereas State-C bout durations were exponentially distributed (Extended Data Fig. 8d) resembling the distribution of sleep bouts across mammalian species^32^. Thus, although the EEG/EMG features of State-C resembled those of light sleep, its episode durations were closer to those of SWS, so that no single sleep stage accounted for the state. Interm-C and State-C thus alternated on short timescales (Fig. 4d), with State-C episodes often briefly interrupted (0.5 to 10 s) by short movements (“shifting”) that coincided with transient EEG γ activation, especially during “corner” postures (Extended Data Fig. 7b; Extended Data Movie 12). These alternating dynamics resemble sleep–wake transitions and are consistent with flip-flop-like arousal-state switching mechanisms involving the ascending reticular activating system (ARAS)^31,33^.

To examine the time course during the active phase, we housed mice on an inverted light–dark cycle. State-C occupied a fraction of time similar to that observed during the normal light phase (31% ± 4.6%, *n* = 49). State-C, impaired laser chasing and delayed feeding initiation appeared by ∼5 min after A/C and resolved by ∼60 min (Fig. 4g; Extended Data Fig. 6), paralleling the transient SynC–A/C blockade of spine enlargement in dissociated culture (Fig.1f; Extended Data Fig. 3b,c). Rotarod performance was normal one day after training even when A/C administered shortly after training had induced State-C (Fig. 2e), indicating preserved consolidation of the newly acquired motor memory. A/C-induced State-C was also reproducible three weeks later in 7 previously responsive mice (Fig. 4e), indicating no evident tolerance.

### Abrupt transitions into State-C

Transitions into State-C occurred with similar probability in the light and dark phases, including during the active phase when mice are normally awake (Fig. 4e,g). The associated cognitive impairments were also reproduced in the dark phase (Fig. 4g), arguing against a purely circadian or sleep-drive explanation and suggesting that widespread cortical SynC activation destabilized arousal across circadian states. To characterize the temporal structure of these transitions, we performed time-resolved analyses of Interm-C→State-C transitions (Fig. 4d; Extended Data Fig. 9a–c). In untreated control mice, spontaneous sleep onset proceeded gradually, with progressive declines in locomotor velocity, EMG tone, and cortical γ-band power over 10 to 60 s (Extended Data Fig. 9a; Extended Data Movie 13). In contrast, SynC@FPC mice after A/C exhibited abrupt transitions, marked by rapid decreases in locomotor velocity and cortical γ-band power within ∼10 s (Extended Data Fig. 9b; Extended Data Movie 14). Time-locked averages revealed highly reproducible transition profiles across animals (Extended Data Fig. 9a–c). Despite variability in awake behaviour, this stereotyped transition is consistent with cortical associative activity contributing to arousal stability, potentially through subcortical arousal systems.

Across three behavioural assays—open-field centre exploration, laser-dot chasing and feeding initiation—SynC–A/C consistently impaired performance. These assays likely engage interdependent processes, including perception, attention, motivation and action selection, rather than separable behavioural domains; we therefore interpret their convergent impairment as evidence of impaired integrated cognitive function during goal-directed behaviour^2,34,35^. Because State-C occurred frequently and Interm-C episodes were temporally fragmented, conventional tasks requiring a stable behavioural state over tens of minutes (e.g., T-maze, novel object recognition) could not be performed or interpreted reliably under these conditions. The emergence of State-C further suggests that the impairment extends beyond goal-directed behaviour to cortically mediated stabilization of wakefulness.

### Reduced cortical coupling with preserved mean activity

We performed two-photon Ca^2+^ imaging in the forelimb area of M1 in head-fixed SynC@FPC mice co-expressing GCaMP7f while the mice ran on a treadmill (Fig. 5a,b). The amplitudes and time courses of Ca^2+^ transients were unchanged after A/C in both somatic transients reflecting action potentials (Fig. 5c; Extended Data Fig. 9d) and spontaneous spine Ca^2+^ transients (Extended Data Fig. 9e), indicating that SynC–A/C did not measurably alter the fast Ca^2+^ signals used here. The preserved somatic transients are consistent with intact voltage-gated conductances and spiking, as also indicated by preserved visual responses in V1 (Extended Data Fig. 4). The preserved local spine Ca^2+^ transients—which are NMDA-receptor dependent in vivo^36^—support intact synaptic NMDA-receptor function, consistent with the direct in vitro measurements (Extended Data Fig. 2g). Mice typically ran at mean velocities of 125 ± 18 mm s^−1^ (mean ± s.e.m., *n* = 10 mice, range 20–230 mm s^−1^), which was not affected by i.p. A/C (exact permutation test, *P* = 0.78). Mice were immobile during ∼30% of the recorded time (Fig. 5d). When immobile, Awake mice maintained a stable posture with their legs positioned to grasp the treadmill, whereas during Interm-C leg positions were more variable and poorly controlled (Extended Data Movies 15 and 16).

**Fig. 5.**
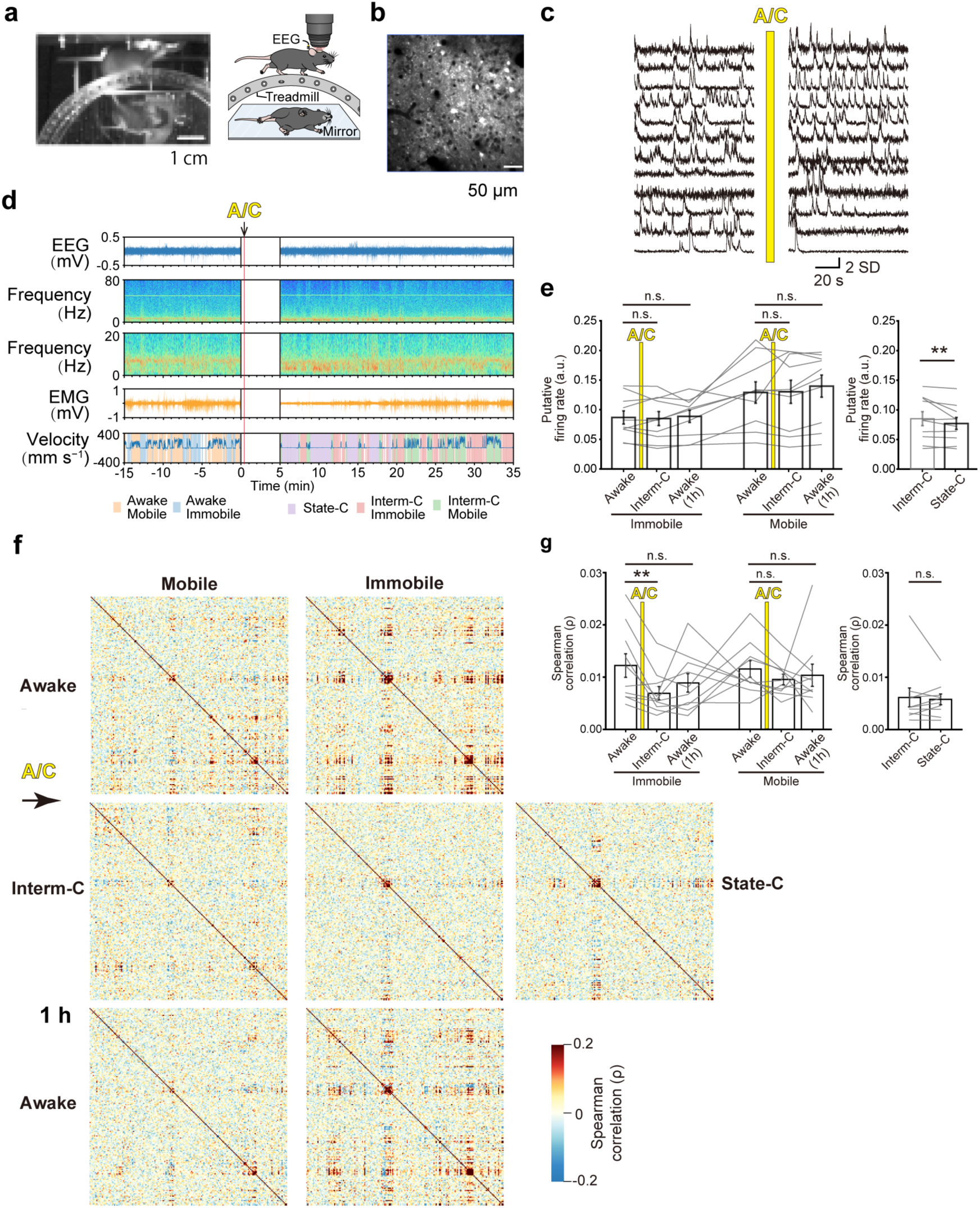
FPC SynC activation reduces M1 ensemble correlations during Interm-C and State-C. Recordings were performed during the animals’ active (dark) phase. **a**, Treadmill setup for two-photon calcium imaging in head-fixed mice. Scale bar, 1 cm. **b**, Example M1 image from a GCaMP7f-expressing SynC@FPC mouse. Scale bar, 50 µm. **c**, Representative GCaMP7f traces from SynC@FPC mice. Scale bars, 2 SD and 20 s. **d**, Simultaneously recorded EEG, its power spectrum, EMG and running speed with behavioural state classification. The gap indicates the interval during which recording was suspended for i.p. injection. **e**, Predicted firing rates derived from Ca^2+^ imaging in 10 SynC@FPC mice, for mobile and immobile phases before i.p. A/C (−45 min to 0 min. Awake), for mobile, immobile Interm-C and State-C phases after A/C (0–45 min), and for mobile and immobile phases 45–120 min after A/C, labelled Awake (1 h). Grey lines, individual mice. Exact permutation test (immobile; mobile in parentheses): Interm-C vs Awake, *P* = 0.63 (0.91); Awake (1 h) vs Awake, *P* = 0.79 (0.35); immobile Interm-C vs State-C, *P* = 0.0020. The Interm-C data are shown again for comparison. The Interm-C vs Awake and Awake (1 h) vs Awake comparisons were tested in an a priori fixed sequence without additional multiplicity adjustment, the second comparison confirmatory only if the first was significant (Methods); the remaining comparisons are secondary and reported unadjusted. **f**, Plots of Spearman correlations for one of 10 SynC@FPC mice during a 100 s period in mobile and immobile states at three periods: Awake, −45–0 min; Interm-C, 0–45 min; and Awake (1 h), 45–120 min. Mobile and immobile phases were separated by running speed (10 mm s^−1^). This example is from a mouse in which correlations returned by 1 h; recovery at 1 h varied across mice (g). **g**, Average Spearman pairwise correlation coefficients in the same 10 mice. Exact permutation test (immobile; mobile in parentheses): Interm-C vs Awake (100 s), *P* = 0.0039 (0.057); Awake (1 h) vs Awake (100 s), *P* = 0.22 (0.68); immobile Interm-C vs State-C (50 s), *P* = 0.89. Fixed-sequence testing as in e. Error bars, s.e.m.

Immobile epochs allowed cortical activity to be compared without a locomotor confound. Firing rates were not significantly reduced after A/C administration during Interm-C (Fig. 5e), but were reduced in State-C, consistent with preserved γ-band power in Interm-C, but not in State-C (Extended Data Fig. 8c). By contrast, pairwise correlations during immobile Interm-C epochs were reduced after A/C (Fig. 5f,g). Concurrently, network dimensionality increased, estimated both by the number of principal components accounting for 50% of the variance and by the participation ratio, each normalized by the number of neurons (D50/N and PR/N; Extended Data Fig. 9f,g)^37^. This combination—reduced pairwise correlations with increased dimensionality—indicates a loss of functional cortical coupling at the level of population activity, rather than generalized suppression of firing. These changes coincided with impaired goal-directed behaviour during Interm-C. Coupling recovered substantially by ∼1 h after A/C, when it no longer differed significantly from baseline (Fig. 5f,g; Extended Data Fig. 9f,g).

State-C epochs identified by EEG and EMG criteria (Fig. 5d) showed a similar pattern of reduced pairwise correlations and elevated dimensionality (Fig. 5f,g; Extended Data Fig. 9f,g), indicating that cortical decoupling extended into low-arousal arrest. We did not attempt a direct imaging-based comparison of State-C with SWS because local somatic Ca^2+^ imaging of pyramidal neurons does not provide a direct readout of the electrophysiological slow-wave synchrony characteristic of SWS^38,39^, and sustained SWS was too rare during treadmill imaging for reliable estimates of pairwise correlation and dimensionality. Behavioural, EEG/EMG and cortical population signatures across Awake, Interm-C, SWS and State-C are summarized in Extended Data Fig. 8e.

### Seconds-scale enlargement of write-permissive neocortical spines is blocked by SynC

To verify the structural target in vivo, we tested whether systemic A/C acutely and reversibly blocked rapid associative spine enlargement in PPC, an optically accessible frontoparietal association area with strongly correlated population activity^40^, using a semi-open cranial window that allowed continuous CDNI-glutamate perfusion and stable two-photon visualization of dendritic spines (Fig. 6a). Pyramidal neurons were sparsely co-labelled with eGFP and the red-shifted optogenetic actuator CsChrimsonR. Three or four dendritic spines within a field of view (FOV) were stimulated by 0.6-ms two-photon glutamate uncaging pulses, followed within 15 ms by a common 2-ms red LED pulse to activate CsChrimsonR and induce postsynaptic depolarization.

**Fig. 6.**
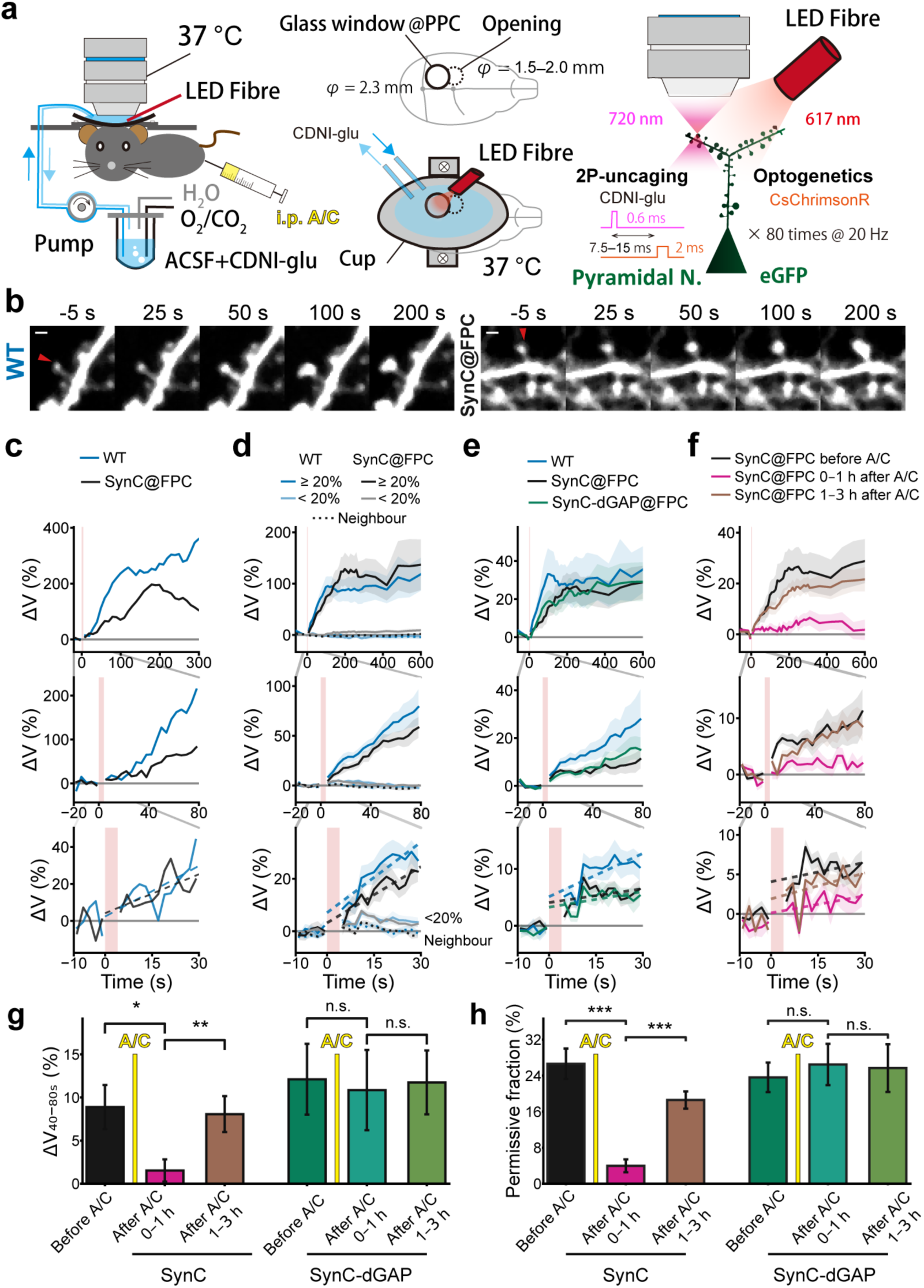
Rapid associative spine enlargement in the posterior parietal cortex (PPC) *in vivo*. **a,** Schematic of the in vivo two-photon uncaging and optogenetics setup. CDNI-glutamate (2.5 mM) in ACSF was continuously superfused through a small circular opening adjacent to the glass window. Cortical pyramidal neurons co-expressed eGFP and CsChrimsonR. A/C was applied i.p., as in the behavioural experiments. The objective lens was heated to 37 °C (Methods). Mice were 4–15 weeks old. **b,** Representative images from spines of a WT mouse (left, 15 weeks old) and a SynC@FPC mouse (right, 6 weeks old) stimulated with glutamate uncaging followed by LED stimulation at 20 Hz for 4 s. Arrowheads indicate the stimulated spines. Scale bars, 1 µm. **c**, Time courses of enlargement (ΔV) of the spines shown in **b**, for 300 s (upper, bin-width 10 s), 80 s (middle, bin-width 4 s) and 30 s (bottom, bin-width 2 s). **d**, Population-average traces for 0–600 s stratified by mean ΔV over 40–80 s (≥20%, dark lines; <20%, pale lines), together with neighbouring-spine traces (dotted), for WT (82 spines, 24 fields of view [FOVs], 2 mice; 21 exceeded 20%) and SynC@FPC before A/C (110 spines, 30 FOVs, 5 mice; 19 exceeded 20%). Dashed lines are linear fits over 4.2–30 s; the 0–4 s stimulation period (pink band) was treated as missing (c–f). Lines and shading, FOV-equal means and s.e.m. **e**, Average ΔV time courses of all stimulated spines in WT and SynC@FPC before A/C (cohorts as in d) and in SynC-dGAP@FPC mice (79 spines, 21 FOVs, 4 mice). **f,** Average ΔV time courses in SynC@FPC mice before A/C (as in d), 0–1 h after A/C (55 spines, 16 FOVs, 5 mice) and 1–3 h after A/C (115 spines, 33 FOVs, 5 mice). **g,** Mean ΔV over 40–80 s after stimulation, showing acute loss of the response and its recovery. SynC@FPC cohorts are those in f and the SynC-dGAP@FPC before-A/C cohort is that in e, with SynC-dGAP@FPC 0–1 h (54 spines, 14 FOVs, 4 mice) and 1–3 h after A/C (70 spines, 20 FOVs, 3 mice). SynC contrasts were assessed using heteroscedastic Normal parametric-bootstrap tests of equally weighted FOV means (Methods). SynC before versus 0–1 h after A/C, *P* = 0.0488; SynC 1–3 h versus 0–1 h after A/C, *P* = 0.0059; SynC-dGAP before vs 0–1 h after A/C, *P* = 0.762, and SynC-dGAP 0–1 h vs 1–3 h after A/C, *P* = 0.481. **h,** Estimated permissive fraction of stimulated spines, showing that the loss in g reflects a change in this fraction. Same cohorts as in g. Bars show the fitted permissive fraction π for each condition, estimated across spines; error bars indicate s.e.m. across FOV-level values, each obtained by averaging posterior permissive probabilities across the stimulated spines of that FOV. Between-condition differences were assessed using parametric-bootstrap tests of FOV means, with the jointly fitted mixture parameters held fixed (Extended Data Fig. 10, Methods): SynC before vs 0–1 h after A/C, *P* < 1 × 10^−4^; SynC 1–3 h vs 0–1 h after A/C, *P* < 1 × 10^−4^; SynC-dGAP before vs 0–1 h after A/C, *P* = 0.585; SynC-dGAP 0–1 h versus 1–3 h after A/C, *P* = 0.904.

This protocol induced rapid spine enlargement in PPC pyramidal neurons, detectable within the initial 0–30-s period, without apparent delay in WT and SynC@FPC spines (Fig. 6b–d): The least square fitting of the spine enlargement between 4.2 and 30 s indicated the onset of spine enlargement within 4 s, or shorter. Enlargement with a similar time course was detected in the average traces of spines with more than 20% enlargement (WT mice, Fig. 6d). In the remaining ∼80% of spines, enlargement was below 20% and, on average, undetectable (Fig. 6d, pale traces). Nevertheless, averaged over all stimulated spines, without stratification, enlargement was still detected in SynC@FPC before A/C and in SynC-dGAP@FPC mice (Fig. 6e), and in SynC@FPC 1–3 h after i.p. A/C, whereas it was nearly abolished 0–1 h after A/C (Fig. 6f,g). Fitting the response distribution (see below) attributed this loss to a fall in the fraction of spines that enlarged, from 26.6% before A/C to 4.1% within 1 h (55 spines; *P* < 1 × 10^−4^; Fig. 6h), rather than to a uniform reduction across all spines. Thus, the rapid spine enlargement was blocked by Rac1-GAP catalytic activity rather than by A/C treatment, SynC expression or FKBP recruitment itself.

To confirm the validity of this conclusion, we analysed the distribution of responses across all 565 stimulated spines, rather than their mean. Mixture-model analysis, in which the permissive component was an exponentially distributed enlargement (θ = 34.4%) observed with the same normal noise as neighbouring spines, identified a write-permissive fraction (π) of approximately 24% of stimulated spines at the time of stimulation, characterized by 40–80-s enlargement exceeding the baseline distribution of neighbouring spines (Extended Data Fig. 10a). Without any fitting, the neighbouring-spine distribution (σ = 9.37%) predicts that only 0.4 of 565 spines would exceed ΔV = 30%, yet 50 did, spread over 44 of 158 FOVs and all 11 mice—close to the 45 FOVs expected if such spines were distributed at random—indicating that write-permissive and write-protected spines coexist (Extended Data Fig. 10b), consistent with uncaging results in V1^41^. This selective write permissiveness accords with the neocortex operating as a slower-learning system than the hippocampus^42^ and may help explain why cortical structural plasticity has been difficult to detect experimentally. It also provides cellular support for the write-protected synaptic architecture proposed in cascade-type memory models^43^, in which rapid updating is balanced against indiscriminate overwriting during continuous activity.

Thus, rapid associative spine enlargement takes place in the very cortical network in which it sustains spike correlations and goal-directed behaviour. How rapid associative spine enlargement—and its restriction to a write-permissive subset—is expressed through coordinated pre-, post- and trans-synaptic functional changes remains an important direction for future work.

## Discussion

We developed SynC, a PSD-targeted (Fig. 1c and 3b; Extended Data Fig. 1b and 5e), A/C-activated synaptic chemogenetic perturbation that acutely and reversibly suppresses activity-dependent spine enlargement (Fig. 1e,f; Extended Data Fig. 3a–c; Fig. 6f,g) while sparing basal spine dynamics (Fig. 1g, Extended Data Fig. 3e), synaptic transmission and NMDA receptor–mediated responses (Extended Data Fig. 2e–g; Extended Data Fig. 9e), intrinsic excitability (Extended Data Fig. 2c,d), short-term plasticity (Extended Data Fig. 2h,i), spine shrinkage (Extended Data Fig. 3d), and basic cortical sensory responses (Extended Data Fig. 4). In vivo two-photon uncaging further revealed seconds-scale associative spine enlargement in neocortical pyramidal neurons and its reversible block by SynC–A/C (Fig. 6). With local M1 expression, SynC–A/C selectively abolished motor learning while sparing initial motor performance (Fig. 2d); with broad frontoparietal expression, SynC–A/C impaired goal-directed behaviours, notably delaying feeding initiation despite preserved locomotion and consummatory behaviour (Fig. 3c–e; Extended Data Fig. 6). Notably, a similar dissociation— spontaneous feeding lost, consummatory responses spared—had classically required interventions as drastic as ablating the cerebral cortex^44,45^ or disconnecting the forebrain^26^; here it appeared with the brain intact, from an acute, reversible synaptic block by SynC–A/C that resolved within an hour. During behaviourally impaired Interm-C, mean firing rates (Fig. 5e) and γ power were preserved (Fig. 4c, Extended Data Fig. 8c), whereas pairwise spike correlations decreased (Fig. 5g), indicating disrupted cortical coordination rather than acute neuronal toxicity or generalized cortical suppression.

The rapid onset and within-hour recovery of these cellular, circuit and behavioural effects argue against developmental, compensatory or irreversible changes. Preserved consolidation of a newly acquired motor memory despite acute SynC–A/C perturbation (Fig. 2e), together with the reproducible re-induction of State-C in the same mice three weeks later (Fig. 4e), provides further evidence against lasting dysfunction or tolerance. Specificity and state-classification controls showed that GAP-dead SynC preserved A/C-driven FKBP recruitment (Extended Data Fig. 1d) but lacked Rac1-GAP catalytic activity (Fig. 1e) and produced no rapid synaptic (Extended Data Fig. 1c–f; Fig. 6g,h), behavioural (Fig. 2d, 3c–e), or arousal phenotype (Fig. 4e). A/C alone and the FKBP-Rac1-GAP-alone (+A/C) control did not induce State-C (Fig. 4e); and the EEG/EMG profile distinguished State-C from anaesthesia, absence-like seizures, cataplexy and other arrest-like states (Supplementary Table 1). Together, these findings and controls support the interpretation that SynC selectively and reversibly suppresses rapid spine enlargement, and that this underlies the circuit and behavioural consequences.

SynC–A/C produced State-C, an arrest state whose EEG/EMG profile shared elements with sleep—reduced EMG tone and spindles—yet lacked the robust slow-wave activity (SWA) and sleep postures that define SWS (Extended Data Fig. 7b,c, 8c; Fig. 4c,d), whereas promoting spine enlargement with SynK–A/C increased SWA^13^. These opposite effects on SWA support the view that SynC acts predominantly by blocking activity-dependent spine enlargement, a structural process closely coupled to synaptic potentiation^6,7,9^. State-C was accompanied by an autonomic shift consistent with reduced arousal—miosis and modest heart-rate slowing without respiratory suppression (Fig. 4f)—and its bout durations were exponentially distributed^32^, consistent with engagement of an endogenous arousal-state switching mechanism (Fig. 4d; Extended Data Fig. 8d). Wakefulness is also known to become less stable under monotonous or low-engagement conditions^46^. Cortical neurons can influence sleep-wake state^47,48^, with extensive corticofugal projections providing a plausible substrate for such influence^49,50^. More broadly, preserved wake-like firing and γ activity alone were insufficient for awake cognitive function. When rapid associative spine enlargement was acutely blocked, cortical activity remained globally active, yet pairwise correlations fell, goal-directed behaviours failed, and wakefulness became unstable. The acute reduction of spike correlations (Fig. 5g) suggests that enlargement is ongoing during wakefulness. Because enlargement is induced by coincident pre- and postsynaptic activity (Fig. 6), enlargement and correlated activity reinforce each other. Thus, seconds-scale associative spine enlargement is an active online synaptic process required to maintain functional cortical coupling, cognitive function and stable wakefulness.

## Methods

### Animals

Cortical neurons for dissociated culture were obtained from embryonic day 17 (E17) ICR mouse embryos (Japan SLC). Sprague–Dawley rats (Sankyo Lab Service) were used to prepare organotypic slice cultures. Neurons from both sexes were mixed in these experiments.

For SynC@FPC experiments, 151 male and 162 female C57BL/6J mice (Sankyo Lab Service) received viral injections at postnatal day 1 (P1). Because sex determination at P1 is impractical, both sexes were included, and data from males and females were pooled since no apparent sex differences were observed (e.g., Fig. 4b). Thy1-GCaMP6s transgenic mice (C57BL/6N background, Japan SLC) were used for Ca^2+^ imaging experiments in visual cortex (V1). Only male C57BL/6J mice aged 8 weeks were used for the rotarod experiments (Fig. 2), and only males for the V1 imaging (Extended Data Fig. 4), to ensure consistency with existing normative datasets and to minimize animal numbers, as these paradigms were not specifically designed or statistically powered to detect sex differences.

Animals were housed under controlled temperature and humidity conditions and typically maintained on a 12-h light/dark cycle. At the time of experiments, animals were 8–17 weeks old, except for the in vivo uncaging experiments (Fig. 6, Extended Data Fig. 10), in which mice were 4–15 weeks old. For experiments shown in Fig. 4, Fig. 5, Extended Data Fig. 6, and Extended Data Fig. 9, the light/dark cycle was reversed to assess behavioural performance during the animals’ active (night) period. All experimental procedures were approved by the Animal Experimental Committee of the Faculty of Medicine, The University of Tokyo (A2024M029-03).

### Plasmid constructions and adeno-associated virus (AAV) preparation

The following plasmids were generated by PCR and In-Fusion cloning (Takara Bio):

- pAAV-CaMKII(0.4)-PSDΔ1,2-FRB-IRES-HA-FKBP-GAP-W (SynC-fusion);
- pAAV-CaMKII(0.4)-PSDÄ1,2-FRB-IRES-HA-FKBP-GAP-dead(R179G)-W (SynC-dGAP-fusion);
- pAAV-CaMKII(0.3)-PSDΔ1,2-FRB-W;
- pAAV-CaMKII(0.3)-PSDΔ1,2-FRB-mVenus-W;
- pAAV-CaMKII(0.3)-mVenus-FKBP-GAP-W;
- pAAV-CaMKII(0.3)-mVenus-FKBP-GAP-dead(R179G)-W;
- pCAG-mScarlet-W;
- pAAV-CaMKII(0.4)-Cre-W;
- pAAV-hSyn-DIO-mScarlet-W;
- pAAV-hSyn-DIO-PSDΔ1,2-FRB×3-W;
- pAAV-hSyn-DIO-mVenus-FKBP-GAP-W;
- pAAV-hSyn-DIO-mVenus-FKBP-GAP-dead(R179G)-W;
- pAAV-hSyn-DIO-mEGFP-W; and
- pAAV-hSyn-DIO-CsChrimsonR-mScarlet-W.

Human FKBP and FRB, and mouse α1-chimerin (Rac1-GAP) cDNAs were cloned from cDNA templates. GAP denotes the Rac1 GTPase-activating protein domain (amino acids 143–322) of α1-chimerin. The GAP-dead mutant was generated by introducing an R179G substitution into α1-chimerin. PSDΔ1,2 was created by deleting amino acids 65–312 from PSD-95. FRB contained the T2098L mutation, as in SynK^13^, to reduce binding to endogenous FRB and mTOR.

For dissociated neuronal cultures (Fig. 1f; Extended Data Figs. 1 and 3), the indicated plasmids were introduced by lipofection. For slice-culture experiments (Fig. 1e; Extended Data Figs. 2) and in vivo experiments (Figs. 2–6; Extended Data Figs. 4–10), the relevant pAAV constructs were packaged into AAV particles.

AAVs were produced and titrated as described previously^13^. Briefly, plasmids encoding the AAV vector, pHelper (Stratagene), and pUCmini-iCAP-PHP.eB (Addgene #103005) were transfected into HEK293 (AAV293) cells (Stratagene). After three days, cells were harvested and AAV particles were purified twice by iodixanol gradient centrifugation. Viral genome titres were determined by quantitative PCR. AAV1-Syn-jGCaMP7f (#104488-AAV1) was purchased from Addgene and used for calcium imaging. SynC-fusion and SynC-dGAP plasmids have been deposited at Addgene (IDs 248572 and 248573).

### Dissociated culture of neocortex

Cortical neurons were dissociated from embryonic day 17 (E17) ICR mouse embryos and plated at 4.5 × 10^4^ cells cm^−2^ on poly-L-lysine-coated glass-bottom dishes (MatTek, #1.5). Cultures were maintained at 37 °C in 5% CO_2_ in Neurobasal medium (Gibco) supplemented with B27 and L-glutamine. At 16 days in vitro (DIV16), cells were transfected with CAG-mScarlet, CaMKII(0.3)-PSDΔ1,2-FRB×3 and CaMKII(0.3)-mVenus-FKBP-GAP at a plasmid ratio of 5:15:1 using Lipofectamine 2000 (Invitrogen). After 3–5 days, dual-colour time-lapse images (mVenus and mScarlet) were acquired every 20 min using a confocal microscope (A1R, Nikon). Translocation was induced by adding the A/C heterodimerizer (Takara Bio; 0.5 mM stock in ethanol) at a final concentration of 4 µM followed by three washes after 10 min. Ethanol (0.05%) was always contained in the medium.

To induce spine enlargement (cStim), neurons were pre-treated with 50 µM D-AP5 (Abcam) for 12 h before stimulation, then the medium was replaced with a solution containing (in mM): 140 NaCl, 5 KCl, 10 D-glucose, 2 CaCl_2_, 10 HEPES-Na, 200 µM glycine, and 20 µM bicuculline (cStim). After 3–10 min, cultures were returned to the original medium. To induce spine shrinkage (Extended Data Fig. 3d), neurons were treated with 20 µM NMDA for 5 min in the continuous presence of 400 nM muscimol^51^. For analysis of intrinsic spine fluctuations (Fig. 1g, Extended Data Fig. 3e), the coefficient of variation (CV) was obtained from the s.d. and mean values before and after application of A/C or vehicle. For histological examination (Extended Data Fig. 1b), cultured cells were washed with phosphate-buffered saline (PBS, pH 7.4) and fixed in 4% paraformaldehyde in PBS for 15 min at room temperature, then stored in PBS at 4 °C. Immunohistochemistry was performed as described in the Histology section.

### Hippocampal slice culture

Transverse hippocampal slices (350 µm thick) were prepared from postnatal day 6–8 Sprague–Dawley male rats and placed on 0.4-µm culture inserts (EMD Millipore). Slices were maintained at 35 °C in 5% CO_2_ in culture medium containing 50% MEM (Invitrogen), 25% HBSS (Invitrogen), 25% horse serum (Invitrogen), and 6.5 g L^−1^ glucose^6^.

AAV mixtures were injected into area CA1 on the day of or the day after slice preparation under an upright microscope to confine viral spread within CA1. The injected AAVs were as follows: pAAV-CaMKII(0.4)-Cre (5.0 × 10^9^ gc mL^−1^); pAAV-hSyn-DIO-mScarlet (2.0 × 10^13^ gc mL^−1^); pAAV-hSyn-DIO-PSDΔ1,2-FRB × 3 (3.0 × 10^13^ gc mL^−1^); pAAV-hSyn-DIO-mVenus-FKBP-GAP (2.0 × 10^12^ gc mL^−1^); and pAAV-hSyn-DIO-mVenus-FKBP-dGAP (2.0 × 10^12^ gc mL^−1^). After 10–12 days in culture, slices were transferred to a recording chamber and superfused with artificial cerebrospinal fluid (ACSF) containing (in mM): 125 NaCl, 2.5 KCl, 2 CaCl_2_, 1 MgCl_2_, 1.25 NaH_2_PO_4_, 26 NaHCO_3_, 20 glucose, and 200 µM Trolox (Sigma-Aldrich), continuously bubbled with 95% O_2_/5% CO_2_. MgCl_2_ was removed from ACSF for the recording of NMDA receptor-mediated currents. The ACSF initially contained the vehicle used for A/C. Twenty minutes before uncaging, it was replaced with ACSF containing A/C at a final concentration of 4 µM. Synaptic transmission was recorded from CA1 pyramidal neurons under whole-cell voltage clamp using patch electrodes (5– 7 MΩ) filled with internal solution containing (in mM): 120 potassium gluconate, 20 KCl, 10 disodium phosphocreatine, 4 ATP (Mg^2+^ salt), 0.3 GTP (Na^+^ salt), and 10 HEPES-KOH (pH 7.3). Evoked synaptic currents were averaged from 10 consecutive stimuli and plotted at each time point (Multiclamp 700B amplifier, Molecular Devices).

Spine enlargement was induced in Mg^2+^-free ACSF containing CDNI-glutamate (1 mM), as described previously^6,18^. Images were acquired at 950 nm through a water-immersion objective (XLPLN25XWMP2, 25×/1.05 NA; Evident). Two-photon uncaging was performed at 720 nm using 600-µs pulses delivered 80 times at 1 Hz. The uncaging spot was positioned approximately 1–2 pixels outside the distal edge of the spine head, and the laser power measured at the front aperture of the objective was adjusted between 4 and 7 mW according to spine depth. Experiments were conducted at 31 °C.

To isolate receptor-specific components, NBQX (10 µM; MedChemExpress) or D-AP5 (50 µM) was applied together with bicuculline (20 µM; Fujifilm) at a holding potential of −70 mV for NMDA- or AMPA-receptor-mediated currents, respectively, whereas GABA-mediated currents were isolated at +30 mV in the presence of D-AP5 and NBQX. For recordings of synaptic facilitation and depression and post-tetanic potentiation, D-AP5 was included in the bath solution, and QX-314 was included in the intracellular solution. Currents were low-pass filtered at 4 kHz and sampled at 20 kHz. Series resistance was ∼20 MΩ (uncompensated), and recordings with resistance >30 MΩ were excluded.

### Mouse AAV and A/C injection

All AAV-mediated transductions were performed using the AAV.PHP.eB capsid. For adult mice, 1 µL of AAV::CaMKII-SynC (1.0 × 10^13^ gc mL^−1^) was pressure-injected using a syringe pump (RWD) at 0.1 µL min^−1^. Injections were made bilaterally into M1 (AP +1.0 mm, ML ±1.3 mm from bregma, DV −0.75 mm from the brain surface; Fig. 2d) and unilaterally into V1 (AP −3.0 mm, ML +2.0 mm from bregma, at a depth of 200–450 µm; Extended Data Fig. 4). Injections were made through borosilicate glass capillaries without filament (G-1; Narishige) pulled on a P-1000 micropipette puller (Sutter Instrument).

For neonatal mice (postnatal day 1), injection pipettes were fabricated from 10 cm quartz glass capillaries (outer diameter 1.0 mm, inner diameter 0.5 mm; no filament; Q100-50-10, Sutter Instrument) using a P-2000 puller (Sutter Instrument) to yield ∼30 µm tip diameters. Each pipette was attached directly to a 25 µL Hamilton syringe (1702 RN) via a compression fitting. Viral injections were performed in 313 pups at postnatal day 1 (P1). Each pup was anaesthetized by brief hypothermia on an ice bed and secured in a custom neonatal head holder (Narishige; designed by Mr. Ooishi). The holder arms were positioned vertically at bregma level along the anterior–posterior axis, 0.3 mm lateral to the midline on both hemispheres. The glass micropipette was advanced perpendicularly through the skull to a depth of 400-500 µm, and 4 µL of AAV solution was injected using a syringe pump (Neurostar). After recovery on a heating pad, pups were returned to their home cage. The procedure required less than 20 min per pup.

The A/C heterodimerizer (Takara Bio) was dissolved in N,N-dimethylacetamide (Fujifilm) at 20 mg mL^−1^ and diluted in propylene glycol (Nacalai Tesque) to 2 mg mL^−1^. Before use, this stock was diluted to 0.4 mg mL^−1^ in diluent (1.2% Polysorbate 80, Fujifilm; 27% PEG-300, Fujifilm)^52^ and injected intraperitoneally at 2 mg kg^−1^ body weight. For Ca^2+^ imaging (Fig. 5 and Extended Data Fig. 9), AAV::SynC-fusion (1.2 × 10^13^ gc mL^−1^) and AAV::syn-jGCaMP7f (2.0 × 10^12^ gc mL^−1^) were co-injected into neonatal pups.

### Measurement of spine enlargement

Three-dimensional reconstructions of dendritic morphology were generated by the summation of z-stacked images separated by 0.3 µm for confocal (Fig. 1; Extended Data Fig. 1, 2, 3) and by 0.5 µm in slice culture experiments (Fig. 1g). Spine-volume changes were estimated from the total fluorescence intensity of the spine head, using mScarlet for the experiments shown in Fig. 1 and Extended Data Figs. 1–3, and eGFP for the in vivo experiments shown in Fig. 6 and Extended Data Fig. 10. Spine-head fluorescence was normalised to reference fluorescence within the imaging field (Fig. 1 and 6; Extended Data Fig. 1, 2, 3 and 10)^6^. For Fig. 6 and Extended Data Fig. 10, FOVs affected by incomplete stimulation, severe drift or focal-plane instability, and spine ROIs with structural damage or insufficient prestimulation signal above background (mean ROI signal ≤1.25 × background), were excluded without reference to response magnitude or experimental condition.

### Vital signs

EEG, EMG and heart rate (HR) signals were recorded in an electrically shielded open field using a preamplifier (CerePlex Direct, Blackrock) connected to the main amplifier via a lightweight cable. The preamplifier, connected via a 6- or 8-pin adaptor, was secured to the skull using dental cement.

At 8–10 weeks of age, mice were implanted with EEG and EMG electrodes under isoflurane anaesthesia. EEG electrodes were placed over the frontal cortex (AP +1.0 mm, ML +1.5 mm from bregma), with the ground electrode attached to the skull over the cerebellum. EMG was recorded from two wire electrodes inserted into the *m. trapezoideu*s. To reduce mechanical noise when the mouse bumped into the wall, the EEG/EMG preamplifier was enclosed in a plastic protective frame.

EEG signals were filtered at 0.3–250 Hz using a four-pole Butterworth filter, sampled at 1 kHz. The signals were fast-Fourier-transformed to obtain power spectra (0.5-Hz resolution; absolute power in µV^2^ [0.5 Hz]^−1^). Heart rate (HR) was monitored via two subcutaneous wire electrodes, one placed anterior to the heart and the other in the *m. pectoralis major*, connected to the EEG preamplifier. Data were displayed online and analysed offline using custom Python scripts.

Respiration, body temperature and pupil size were monitored in head-fixed mice on a custom turntable (Thorlabs). Running speed was measured using a rotary encoder (Omron). Respiration was detected with a subcutaneous thermocouple electrode placed beneath the nose, and temperature fluctuations were converted to respiratory traces (PTW-401, Unique Medical). Posture and pupil size were video-recorded and quantified using DeepLabCut^53^ (v2.3.10) with custom Python (3.9.13) routines.

### Behaviours

#### Motor learning

For motor learning (Fig. 2), mice were trained on a rotarod (MK-610K, Muromachi). Before training, mice were habituated to the stationary rod until calm for ∼2 min. During training, the rod accelerated linearly from 0 to 80 rpm over 300 s. Each mouse underwent three 5-min on the training day with 1-min inter-trial intervals, and latency to fall was recorded for each trial. Mice were considered to have fallen when they clung to the rod and passively rotated for three consecutive turns; trials in which mice fell while facing the direction of rod rotation were discarded and repeated. Motor learning consolidation was tested on the following day with one accelerating trial (0–80 rpm). The three training trials and the consolidation test were treated as four independent time points, and motor learning was quantified as the area under the resulting learning curve, using the latency of the first trial as the baseline.

#### Open-field tasks

Open-field tests were performed either in a large arena (35 × 35 cm) to measure running velocity (Fig. 3) or in a smaller arena (19.4 × 19.4 cm) for close-up videography of laser chasing, food-pellet feeding, and State-C behaviours (Extended Data Movie 1–16). In both arenas, the behaviour was video-recorded and the position of the mouse was tracked with DeepLabCut; running velocity and time spent in the centre (Fig. 3c) were computed from the tracked position in the large arena. In the large arena, postures were classified by direct visual inspection of the video to obtain the posture composition shown in Extended Data Fig. 7d. Body position and orientation were computed from DeepLabCut-tracked body points and used to distinguish State-C from SWS (Extended Data Fig. 7c; see ***State classification***). The small arena permitted close-up recording and was used for the posture images in Extended Data Fig. 7a,b.

For the **laser-dot task**, a green laser pointer (0.5 mW; Sanwa) was projected onto a corner of the small open field for 10 s with 10-s intervals, repeated four times. The laser dot was kept stationary, and an approach was defined as the mouse moving to the dot until the dot fell on its head. A mouse was scored as responsive if it approached the dot in at least one of the four presentations, thereby excluding the possibility that the mouse simply failed to attend to the stimulus. The laser was projected onto a corner of the arena to avoid spurious reflections from the floor. For the **food-pellet task**, a regular food pellet was dropped into the arena, and the latency from pellet introduction to the first bite while grasping the pellet was measured. Approach and sniffing without grasping were not counted as the endpoint. State-C was identified from posture and EEG/EMG. Time spent in State-C was subtracted from the latencies reported in Fig. 3e and 4g, whereas data from mice that entered State-C were excluded from the time courses in Extended Data Fig. 6. These tasks were quantified either by visual inspection or, for the distance from the mouse to the laser dot or the food pellet, with DeepLabCut (Extended Data Fig. 6). The mice showing prolonged State-C episodes were excluded entirely.

#### State classification

Here, SWS denotes the high-delta, low-EMG sleep state used as the physiological comparator; NREM is used only when referring to conventional sleep-stage classification. **State-C** episodes were identified as a sudden behavioural arrest lasting ≥10 s in non-sleeping posture (typically standing, with the tail not curled around the body) after A/C in SynC@FPC mice (Extended Data Fig. 7d). Because State-C was invariably accompanied by a marked reduction in EEG γ-band power and EMG amplitude,EEG and EMG recordings were also used to support state scoring and resolve ambiguous immobile periods during Ca^2+^ imaging on the treadmill. **Interm-C** epochs were defined as post-A/C periods outside State-C before behavioural recovery. For quiet-immobility analyses, epochs were selected by locomotor immobility after excluding State-C and conventional sleep. For the criteria distinguishing Interm-C and State-C from sleep, seizure, anaesthesia and other arrest-like states, see Supplementary Table 1.

### Cranial window surgery

To construct a cranial window for imaging, mice were anaesthetized with isoflurane (3% for induction, 1% for maintenance) and were secured on a stereotaxic frame (Narishige, Japan). The body temperature was maintained at 36–37 °C with a heating pad. To reduce brain edema, 20% mannitol (2-3 g/kg) was injected intraperitoneally before the craniotomy.

Craniotomies were made using stainless trephine drills (Fine Science Tools, CA, USA) under continuous gentle perfusion with oxygenated ACSF, at the following sites: M1 (at the level of bregma, 1.3 mm lateral; ϕ 2.3 mm), V1 (1 mm anterior and 3 mm lateral to lambda; ϕ 4.0 mm in the left hemisphere for wide-field imaging, ϕ 2.3 mm for two-photon imaging), and PPC (2 mm posterior and 1.3 mm lateral to bregma; ϕ 2.3 mm)^54^. For PPC imaging, the craniotomy was extended manually with a round steel bur (ϕ 0.6 mm; Busch, Germany) to create a contiguous opening (approximately 1.5-2 mm in diameter) for subsequent superfusion of CDNI-glutamate (see below). The dura mater was then carefully removed with fine forceps in PPC preparations only.

Cranial windows were then sealed with coverslips (Matsunami Glass, Osaka, Japan). For M1, a "glass-plug" configuration^52^ was used, consisting of two stacked circular coverslips (No. 1, 0.12–0.17 mm thickness, ϕ 3.0 mm; and No. 3, 0.25–0.35 mm thickness, ϕ 2.0 mm) glued together with UV-curing optical adhesive (NOA81, Norland Products, NJ, USA)^55^; the lower coverslip was inserted into the craniotomy so that the upper coverslip rested on the surrounding skull. For V1 and PPC two-photon imaging, a single coverslip (No. 0, ϕ 2.3 mm) was used, and for V1 wide-field imaging, a single coverslip (No. 0, ϕ 4.0 mm). In PPC preparations, the coverslip covered only the imaging window, leaving the extended opening exposed.

Coverslip edges were sealed with dental cement (RelyX Unicem 2 Clicker, 3M, MN, USA). A custom head-plate was then attached to the skull: a stainless-steel plate (Misumi, Tokyo, Japan) for M1 and V1 wide-field imaging, and a 3D-printed head-plate (DMM.make, Tokyo, Japan) for V1 and a head-plate with chamber (1.2 mL) for PPC two-photon imaging (Fig. 6a). Head-plates were secured with the same dental cement, except for V1 wide-field imaging, in which Super-Bond (Sun Medical, Shiga, Japan) was used. For M1 imaging, EEG and EMG electrodes were implanted with a 4-pin adaptor in the same surgery.

### Visual cortex imaging

For V1 macro imaging, male mice were anaesthetized with isoflurane (5% for induction, 1.0–1.5% for maintenance), and a midline scalp incision was made. Before imaging, chlorprothixene (1.0 mg kg^−1^; Sigma-Aldrich) was administered intramuscularly, and isoflurane concentration was reduced to 0.2–0.5%.

Wide-field calcium imaging was performed using a macro zoom microscope (MVX10, Evident) equipped with a 2 × objective (NA 0.25, MVX Plan Apochromat). GCaMP6s fluorescence was excited with a mercury lamp through a GFP mirror unit (U-MGFPHQ/XL; excitation 488 nm, emission 507 nm; Evident). Fluorescence images (4 × 4 mm field of view; 512 × 512 pixels) were acquired at 5 Hz using an sCMOS camera (Zyla 4.2, Andor). Drifting grating stimuli (40° diameter) were presented at three horizontal positions on a 32-inch LCD monitor (20 cm from the right eye). For each trial, baseline fluorescence (F) was defined as the mean fluorescence during the 1-s period before stimulus onset. The visual response (ΔF) was defined as the mean fluorescence during the stimulus period, and ΔF/F maps were averaged across trials to generate response maps. Imaging data were analysed using custom MATLAB scripts (MathWorks, R2024b).

For SynC@V1 mice, in vivo two-photon calcium imaging was conducted at 8 weeks of age using a two-photon microscope (AXR-MP, Nikon, Japan) equipped with a 20 × objective lens (CFI75 Apochromat LWD 20XC W, Nikon) and a femtosecond laser (Insight DS+; Spectra-Physics) at 950 nm to excite GCaMP6s. Dendritic spines of GCaMP6s-expressing cells were imaged 60–120 µm below the cortical surface. Series of 500–2000 xy images (8 × digital zoom, 512 × 512 pixels) were captured before and after A/C application^56^.

### In vivo two-photon Ca^2+^ imaging of M1

A custom treadmill system was used to monitor locomotion and posture during two-photon Ca^2+^ imaging of M1 activity^55,57^. The treadmill wheel was mounted on translation stages allowing movement along the caudal–rostral and medial–lateral axes, and the head-fixation height was adjustable. A 45° mirror inside the wheel enabled high-speed videography (Basler acA1920-40um) of simultaneous ventral and lateral views. Wheel rotation was monitored by a rotary encoder (E6B2-CWZ3E, 100 P/R; Omron), and pulses were recorded with CerePlex Direct (Blackrock). Wheel angular velocity was recorded at 2 kHz and synchronized with imaging via TTL pulses.

Two-photon excitation imaging of M1 in SynC@FPC mice was performed with a tunable Ti:sapphire laser at an excitation wavelength of 950 nm, using an objective (XLPLN25XWMP2, 25×/1.05 NA, Evident), and a resonant scanner (FV-1000, Evident). The treadmill was positioned beneath the objective (Fig. 5a). Images were acquired at 30 Hz from the forelimb region of M1 (AP +0 mm, ML +0.5 mm from bregma). Laser power measured at the front aperture of the objective did not exceed 50 mW, and imaging periods per mouse were limited to 150 minutes to avoid photodamage.

### Processing and analysis of two-photon Ca^2+^ imaging data

Regions of interest (ROIs) corresponding to neuronal somata (20–180 per field of view) were extracted with Suite2p^58^ (v0.14.5), which also performed non-rigid motion correction, ROI detection/classification, neuropil subtraction and OASIS deconvolution^59^ (τ = 400 ms). Photobleaching was corrected per ROI by single-exponential fitting to low-activity segments. For analyses of putative firing rates, ΔF/F_0_ was computed and the deconvolved traces were taken as firing rates (Fig. 5e). Deconvolved calcium activity was averaged for each cell across all imaging frames belonging to mobile or immobile periods, which were classified based on treadmill rotation speed at the threshold of 10 mm s^−1^.

Correlation and dimensionality analyses were performed using bleach-corrected ΔF/F_0_ without deconvolution for the three periods, −45 to 0 min (Awake), 0 to 45 min (Interm-C) and 45 to 120 min (Awake [1h]) after i.p. A/C. Fluorescence traces were averaged in three-frame bins before further analysis. For mobile and immobile states, 20 non-overlapping segments of 150 consecutive frames (5 s at 30 Hz; 100 s in total) were sampled at random. The segment length and number were chosen so that an equal amount of data was obtained from every mouse and every period. For the comparison between Interm-C and State-C, 10 segments per condition (50 s in total) were used, as the duration of State-C was limited in some mice. For each mouse and sampled segment, pairwise correlations among all recorded cells were calculated using Spearman’s correlation coefficient (*ρ*). Correlation coefficients were Fisher *z*-transformed, averaged across cell pairs and segments, and then transformed back to *ρ*. Population dimensionality was quantified using the number of principal components required to explain 50% of the variance and the participation ratio of the covariance matrix. Both measures were calculated using all cells recorded from each mouse and normalised by the number of cells.

Two-photon Ca^2+^ imaging was also performed in individual dendrites in M1. Dendritic spine Ca^2+^ transients were separately measured from shaft Ca^2+^ transients to isolate NMDAR dependent components (Extended Data Fig. 9d,e).

### Imaging of two-photon glutamate uncaging in PPC in vivo

We used the pup injection of AAV for either SynC or SynC-dGAP together with AAV for CaMKII-Cre and floxed-eGFP and floxed-CsChrimsonR. Mice were implanted with a cranial window over PPC as described above. ACSF containing CDNI-glutamate at a nominal bath concentration of 2.5 mM was recirculated through the PPC chamber at 0.2 mL min^−1^ (Fig. 6a). The closed circulation loop was used to superfuse the solution, which was bubbled with 95% oxygen and 5% CO_2_. The objective lens (XLPLN25XWMP2) was heated to 37 °C to warm up the bathing solution to the same temperature. eGFP-labelled dendrites were excited at 920 nm and imaged as 320 × 320-pixel fields (132.6 × 132.6 µm; 0.414 µm pixel^−1^) at 1.79 frames s^−1^. Images were acquired continuously from approximately −20 to 320 s relative to stimulation, with additional 20-frame image series acquired at approximately −60 s and at 60-s intervals from 360 to 600 s. Imaging laser power measured at the front aperture of the objective was 4–8 mW. The CDNI-glutamate concentration beneath the centre of the 2.3-mm-diameter coverslip, at a depth of 100 µm below the dural surface, was estimated from Alexa Fluor 594 fluorescence to be approximately 40% of the bath concentration (∼1 mM versus 2.5 mM in the bath). Mice were supplied with humidified oxygen gas and warmed to 37 °C with a heating pad (FST-HPS; Fine Science Tools Inc., North Vancouver, Canada) during all surgical and imaging procedures. The mouse was secured to the microscope stage through the chamber with screws, and weakly anaesthetized with 0.5% isoflurane during the imaging period. Dendritic spines were imaged 20–50 µm below the pial surface, and the uncaging laser power measured at the front aperture of the objective was 14–18 mW. Red light (617 nm) was delivered through a 0.4-mm-core optical fibre whose tip was fixed 500 µm above the pial surface at an angle of 20°. Optical power measured at the fibre tip was 5–8 mW. Within each FOV, three or four spines were targeted. During each 50-ms cycle, a 0.6-ms uncaging pulse at 720 nm was delivered to each spine 7.5–15 ms before the common red-LED pulse (2-ms duration). This sequence was repeated 80 times at 20 Hz for 4 s. A/C heterodimerizer was applied intraperitoneally at the same dose as in the behavioural experiments, from a stock diluted twofold with ACSF.

### Histology

Three to ten weeks after behavioural experiments, mice were deeply anaesthetized with 3% isoflurane and perfused transcardially with PBS (pH 7.4), followed by 4% paraformaldehyde (PFA) in PBS. Brains were post-fixed overnight in 4% PFA at 4 °C and then stored in PBS.

For HA-tagged probes, free-floating sagittal sections (50 µm) were cut at 0.5, 1.35, 2.35 and 3.55 mm from the midline using a vibratome (VT1200, Leica). Sections were rinsed in PBS, blocked for 1 h at room temperature (RT) in 5% normal goat serum (NGS) in PBST (PBS containing 0.2% Triton X-100), incubated in primary antibody solution (5% normal serum, 0.2% Triton X-100 in PBS) for 24 h at 4 °C, and then in secondary antibody solution under the same conditions for 2 h at RT. To detect translocation of the HA-tagged probe, spines whose HA labelling exceeded that of the parent dendritic shaft were scored as HA-positive in thin dendritic regions, which lie mainly in layer 1. Such spines were seen after A/C but not before (Fig. 2b).

#### Primary antibodies

- Mouse anti-HA (Alexa Fluor 488-conjugated, clone 6E2, #2350S; Cell Signaling Technology), 1:200
- Mouse anti-HA (Alexa Fluor 647-conjugated, clone 6E2, #3444S; Cell Signaling Technology), 1:200
- Mouse anti-HA (clone 6E2, #2367S; Cell Signaling Technology), 1:200
- Rabbit anti-NeuN (EPR12753, ab177487; Abcam), 1:400
- Mouse anti-phospho-neurofilament-H (SMI-31, NE1022; Sigma-Aldrich), 1:1000
- Mouse anti-PSD95 (7E3-1B8, MA1-046; Thermo Fisher), 1:500
- Mouse anti-Homer 1/b/c (SY-72G2; Synaptic Systems), 1:400

#### Secondary antibodies

- Goat anti-rabbit IgG (Alexa Fluor 594, #A-11012; Thermo Fisher), 1:500
- Goat anti-rabbit IgG (Alexa Fluor 405, #A-31556; Thermo Fisher), 1:500
- Goat anti-mouse IgG1 (Alexa Fluor 594, #115-587-185; Jackson ImmunoResearch), 1:500

Nuclei were counterstained with DAPI (4′,6-diamidino-2-phenylindole; ab228549, Abcam) and mounted using Vectashield Antifade Mounting Medium (Vector Laboratories, Burlingame, CA, USA), then cover-slipped.

For mVenus-labelled SynC probes (without HA tag), HA staining was omitted. Sections were rinsed in PBS and directly mounted. Whole-brain serial sections were imaged with a confocal microscope (A1R, Nikon and FV-5000, Evident). Z-stack images acquired at 4 µm steps for sagittal sections (8 µm total) and at 0.4 µm steps for higher-magnification views (17.6 µm total), then aligned and collapsed using NIS-Elements Advanced Research software (Nikon). Histological verification of SynC expression was obtained in 24 of the 42 mice used for behavioural experiments (21 of 32 mice contributing to Fig. 3 and 10 of 12 contributing to Fig. 4g).

Imaging and histological analyses were performed using Fiji/ImageJ (v1.54p). SynC expression levels were quantified as Z-scores relative to the s.d. of unlabelled regions and plotted on the Swanson flat map (Extended Data Fig. 5).

Fluorescence images were displayed using linear intensity scaling. Where display levels were adjusted, only linear changes in brightness and contrast were applied; no nonlinear intensity transformations were used.

### Statistics and reproducibility

Data are reported as mean ± s.e.m. with individual data points shown wherever possible. Throughout, *n* denotes the number of biological replicates — mice, cells, dendrites or spines as specified in each figure legend — with the number of independent animals or cultures given alongside. Because most datasets were small and not assumed to follow a normal distribution, non-parametric tests were used by default: two independent groups were compared with the Mann–Whitney *U* test, and paired or repeated measurements within the same unit with the Wilcoxon signed-rank test; comparisons among more than two groups used the Kruskal–Wallis test followed by Steel’s test against a common control, or, for repeated measures, the Friedman test followed by Wilcoxon signed-rank tests; proportions (for example, fractions of write-permissive spines) were compared with Fisher’s exact test, and pairwise neuronal co-activity with Spearman’s rank correlation. Measures that met normality assumptions — intrinsic membrane properties and the coefficient of variation of spine volume — were analysed with repeated-measures or one-way ANOVA and t-tests. All tests were two-sided unless otherwise stated, and where multiple comparisons were made, *P* values were corrected by the Bonferroni method unless otherwise specified; Steel’s test intrinsically accounts for comparisons against a shared control. Exact *P* values, test statistics and degrees of freedom are given in the figure legends. In the figures, asterisks denote \**P* < 0.05, \*\**P* < 0.01 and \*\*\**P* < 0.001; n.s., not significant.

For paired comparisons in Fig. 5e,g and Extended Data Fig. 9f, we used exact permutation tests with the mean within-mouse difference as the test statistic, enumerating all sign assignments. After-versus-Before and 1-h-after-versus-Before comparisons were tested in that prespecified fixed sequence, with the second considered confirmatory only if the first was significant^60^. No additional multiplicity adjustment was applied within the sequence.

Stimulated-spine enlargement (Fig. 6g) was quantified as the mean percentage change in spine volume over 40–80 s after stimulation. Spine responses were first averaged within each FOV, and equally weighted FOV means were treated as the analysis observations. Group differences were assessed using a heteroscedastic Normal parametric bootstrap with group-specific FOV-level residual variances (10,000 replicates per reported contrast). The SynC before-versus-0–1 h after A/C contrast was designated as the primary comparison, and the SynC 1–3 h-versus-0–1 h recovery contrast as secondary; both were reported without multiplicity adjustment. The dGAP contrasts were descriptive and unadjusted. To estimate the permissive fraction (Fig. 6h; Extended Data Fig. 10), stimulated-spine responses were fitted with a mixture of a zero-centred Normal distribution estimated from all pooled neighbouring spines and a non-negative Exponential component convolved with the Normal distribution. The Normal standard deviation and Exponential scale were shared across conditions, whereas the mixture fraction was condition-specific. Posterior permissive probabilities were calculated for individual spines and averaged within each FOV. Between-condition comparisons of FOV-mean posterior permissive probabilities used the same parametric-bootstrap framework, with the jointly fitted mixture parameters held fixed during resampling.

No statistical methods were used to predetermine sample size; sample sizes were comparable to those reported in previous studies of spine plasticity and cortical-state dynamics. Data exclusions were minimal and pre-established: neonatal SynC-injected mice that were unhealthy owing to insufficient maternal care were excluded, and in Fig. 3c–e mice that did not exhibit State-C were excluded because their cortical SynC expression was insufficient (Fig. 4b). For State-C scoring, episodes were excluded only when the video view did not allow the criteria to be applied. Blinding was not feasible for the behavioural experiments, because the A/C-induced behavioural arrest of SynC@FPC mice makes group identity apparent to the observer. To limit observer bias, the distance and velocity measures in Fig. 4g and Extended Data Fig. 6 were extracted automatically with DeepLabCut. Histological quantification was performed by an experimenter who was not involved in the design of the study and was unaware of the group assignment. Data collection and analysis were otherwise not blinded. No formal randomization was performed; however, P1 viral injections were carried out across multiple litters and days without selection for sex, litter, or injection order, effectively randomizing animals across experimental groups. Key findings were reproduced independently: the rotarod and treadmill experiments were performed by two experimenters and reproduced in two laboratories, and State-C was reproduced in three laboratories. Open-field experiments were conducted mainly by one experienced experimenter, who reproduced the State-C phenotype in more than two independent sessions separated by three weeks.

### Use of large language models

GPT-5 (OpenAI) and Claude Opus 5 (Anthropic) were used during manuscript preparation for English editing and for checking internal consistency.

## Acknowledgements

We thank S. Fujii, C. Fujinami, and Y. Hama for technical assistance, R. Hira for introducing the neonatal (pup) injection method and for discussions, Xu, K. Aihara, T. R. Hira, T. Toyoizumi, K. Yoshida, Y. A. Iriki, T. Yamamori, Y. Hayashi, M. Kashiwagi, N. Kitajima, N. Matsumoto and M. Miyano for helpful discussions, during manuscript preparation.

## Author contributions

All experiments were organized by H.K. T.S. led the team (S.Z., H.O., T.A., M.O. and S.K.P.) for synapse-tool design, validation and behavioural application. H.K. and S.Y. provided guidance. T.H., and K.O. led visual-cortical experiments. S.T., M.K. and M.M. contributed to motor-cortical experiments. The manuscript and figures were prepared jointly by all authors.

## Data availability

All data are available in the main text or the Supplementary Materials. DNA plasmids of SYNCit-C (SynC-fusion) and SYNCit-dGAP (SynC-dGAP-fusion) are deposited at Addgene (ID 248572 and 248573). Code availability: Code for the electrophysiological, imaging, behavioural and statistical analyses, together with the model simulations, is available at https://github.com/HaruoKasai/SynC-paper-site. A browsable interface is provided at https://sync-paper-analysis.hkasaimd.chatgpt.site/.

## Materials & Correspondence

Haruo Kasai

## Figures

**Extended Data Fig. 1.**
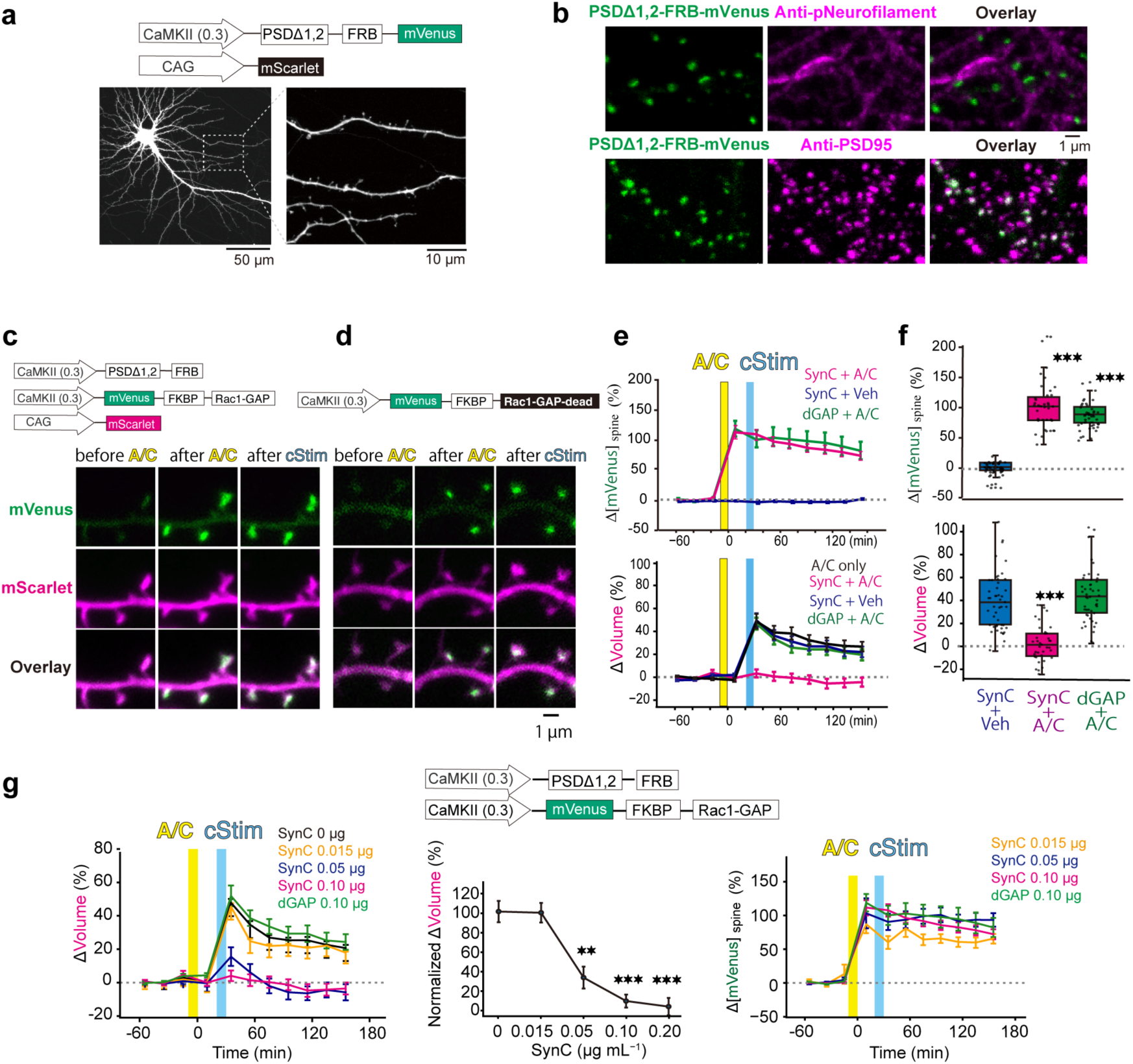
Validation and specificity of SynC in dissociated neocortical cultures. **a**, Normal morphology of dissociated mouse cortical neurons visualized with mScarlet, expressing SynC split components depicted above (1.5 µg mL^−1^ PSDΔ1,2-FRB-mVenus and 0.5 µg mL^−1^ CAG-mScarlet). The right image magnifies the dashed area on the left image. Scale bars, 50 µm and 10 µm. **b**, Subcellular specificity of PSDΔ1,2-FRB-mVenus (green, 3 µg mL^−1^). PSDΔ1,2-FRB-mVenus colocalizes with the postsynaptic marker PSD95 (bottom) and is excluded from axons labelled by phospho-neurofilament (top). Fluorescence images are displayed with linear intensity scaling throughout. Scale bar, 1 µm. **c**, Top, SynC component constructs under the CaMKII(0.3) promoter. Bottom, representative images of mVenus-FKBP-Rac1-GAP (0.1 µg mL^−1^, green) and a volume marker (0.5 µg mL^−1^, mScarlet, magenta). A/C application (yellow) induces rapid enrichment of mVenus at spines, followed by cStim induction (blue). **d**, GAP-dead control. The GAP-dead mutant (0.1 µg mL^−1^) translocates to spines after A/C application but does not prevent cStim-induced spine enlargement. Scale bar, 1 µm. **e**, Time course of spine mVenus enrichment (mVenus fluorescence normalized by the spine volume marker; top) and normalized spine-head volume (bottom). Conditions: SynC + A/C (magenta, *n* = 19 dendrites, 151 spines, 3 cultures), SynC + vehicle (blue, *n* = 26 dendrites, 210 spines, 3 cultures), SynC-dGAP + A/C (green, *n* = 26 dendrites, 237 spines, 4 cultures), and A/C only (black, *n* = 23 dendrites, 196 spines, 3 cultures). **f,** Summary box plots of spine mVenus enrichment (top) and spine-head volume change (bottom) at 20 min post-cStim (same conditions as in **e**). Boxes show median and interquartile range; whiskers show min–max. Kruskal–Wallis, *H*(2) = 93.18, *P* = 5.85 × 10^−21^; Steel’s post hoc vs SynC + vehicle, *P* = 1.28 × 10^−14^ for SynC + A/C; *P* = 3.3 × 10^−16^ for SynC-dGAP + A/C. Enlargement: Kruskal–Wallis, *H*(2) = 58.88, *P* = 1.63 × 10^−13^; Steel’s post hoc vs SynC + vehicle, *P* = 1.38 × 10^−10^ for SynC + A/C; *P* = 0.607 for SynC-dGAP + A/C. **g**, Dose dependence of SynC split (Top). Average blockade of spine enlargement (left), dose dependence of inhibition of spine enlargement (middle), and translocation of FKBP-mVenus-GAP into spines (right) in cells 20 min after A/C application. Cells were lipofected with 1.5 µg mL^−1^ PSDΔ1,2-FRB plasmid (SynC-split). *n* = 23 dendrites (196 spines, 7 cells, 3 cultures) without FKBP-GAP (control); *n* = 17 dendrites (159 spines, 8 cells, 2 cultures) with 0.2 µg mL^−1^ FKBP-GAP; *n* = 19 dendrites (151 spines, 6 cells, 3 cultures) with 0.1 µg mL^−1^; *n* = 17 dendrites (163 spines, 7 cells, 2 cultures) with 0.05 µg mL^−1^; *n* = 21 dendrites (164 spines, 7 cells, 3 cultures) with 0.015 µg mL^−1^; *n* = 26 dendrites (237 spines, 10 cells, 4 cultures) with FKBP-dGAP (SynC-dGAP). Dose-dependent inhibition was tested with Kruskal–Wallis test (*H* = 67.68, *P* = 3.12 × 10^−13^) followed by Steel multiple comparisons vs control: *P* = 8.4 × 10^−7^ for 0.2 µg mL^−1^ FKBP-GAP; *P* = 5.00 × 10^−6^ for 0.1 µg mL^−1^; *P* = 3.46 × 10^−3^ for 0.05 µg mL^−1^ FKBP-GAP; *P* = 1.00 for 0.015 µg mL^−1^ FKBP-GAP; *P* = 1.00 for 0.10 µg mL^−1^ FKBP-dGAP. Relative translocation of FKBP-GAP quantified by ΔmVenus fluorescence normalized by mScarlet fluorescence of each spine (Δ[mVenus]_spine_) was uncorrelated with FKBP-GAP concentration, as PSDΔ1,2-FRB concentration (1.5 µg mL^−1^) was fixed.

**Extended Data Fig. 2.**
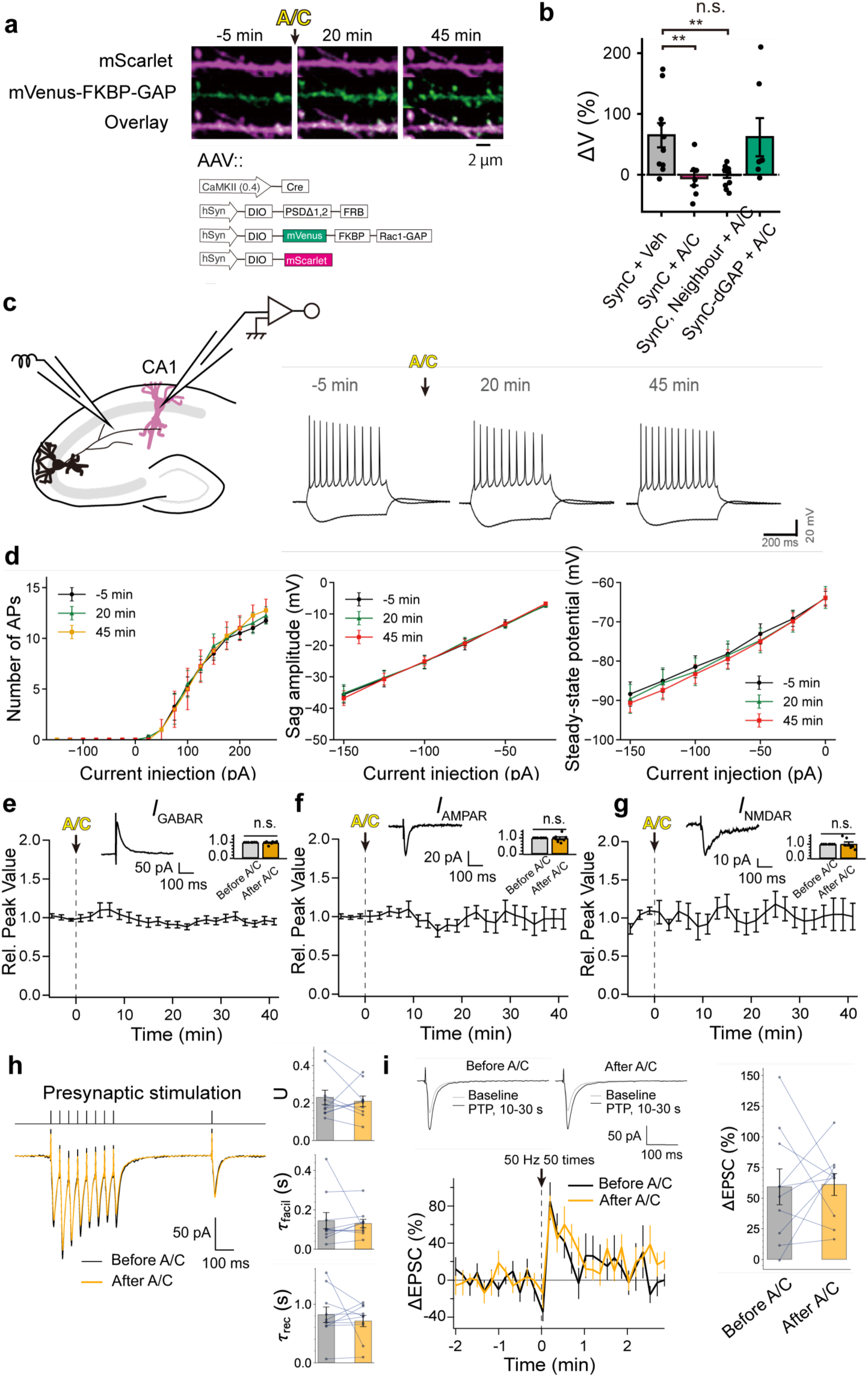
SynC blocks uncaging-induced spine enlargement while sparing neuronal excitability and synaptic transmission in hippocampal slice cultures. **a**, CA1 pyramidal neurons were transduced with AAVs encoding SynC constructs (bottom) for 10-12 days (Methods). Rows show the filler (mScarlet, top), mVenus-FKBP-GAP (middle) and the merged image (bottom); columns show the times relative to A/C application. Scale bar, 2 µm. **b**, Statistical analysis of the uncaging-induced spine-enlargement experiment shown in Fig. 1e. Spines were stimulated 20 min after A/C application, and ΔV (%) was quantified as the mean response 30–40 min after uncaging. Kruskal–Wallis test: *H*(3) = 14.91, *P* = 1.89 × 10^−3^. Pairwise Mann–Whitney *U* tests vs SynC + vehicle, with Holm adjustment: SynC + vehicle (*n* = 10 spines, 2 slices, 2 rats); SynC + A/C, *P* = 9.26 × 10^−3^ (*n* = 7 spines, 2 slices, 2 rats); SynC neighbour + A/C, *P* = 8.09 × 10^−3^ (*n* = 12 spines, 2 slices, 2 rats); and SynC-dGAP + A/C, *P* = 0.813 (*n* = 7 spines, 2 slices, 2 rats). **c**, Schematic of the slice-recording configuration for current-evoked action potentials (APs) and electrically evoked synaptic currents. Representative AP trains are shown before and after A/C (times indicated). **d**, Intrinsic membrane properties were stable across A/C administration. Summary plots show the number of APs vs injected current (left), sag amplitude (middle), and steady-state membrane potential (right) (*n* = 4 neurons, 4 slices, 2 rats). Repeated-measures ANOVA: number of APs, *F*(2,6) = 0.096, *P* = 0.91; sag amplitude, *F*(2,6) = 1.22, *P* = 0.36; steady-state potential, *F*(2,6) = 1.34, *P* = 0.33. **e***–***g**, Synaptic currents recorded in the presence of selective blockers (Methods). A/C had no detectable effect on GABA_A_ receptor–mediated currents (**e**; I_GABAR_), AMPA receptor–mediated currents (**f**; I_AMPAR_), or NMDA receptor–mediated currents (**g**; I_NMDAR_) (*n* = 7 cells each, 7 slices, 3 rats). Wilcoxon signed-rank tests: I_GABAR_, *W* = 12, *P* = 0.81; I_AMPAR_, *W* = 12, *P* = 0.81; I_NMDAR_, *W* = 13, *P* = 0.94. Scale bars, 200 ms and 20 mV (AP traces in c), and as indicated for synaptic-current traces (**e**–**g**). Data are mean ± s.e.m. **h**, Synaptic facilitation and depression studied with a train of eight presynaptic stimuli followed by a single recovery stimulus, delivered through a glass electrode to SynC-transduced cells. A/C did not induce significant changes in the parameters of short-term plasticity, U, τ_facil_ and τ_rec_ (*n* = 10 cells, 10 slices, 3 rats)^21^. Wilcoxon signed-rank tests: U, *W* = 20, *P* = 0.49; τ_facil_, *W* = 17, *P* = 0.32; τ_rec_, *W* = 26, *P* = 0.92. **i**, Post-tetanic potentiation (PTP) induced by 50 presynaptic stimuli at 50 Hz (*n* = 10 cells, 10 slices, 3 rats) in the presence of D-AP5 (50 µM), with QX-314 (1 mM) included in the intracellular solution. Both decays and amplitudes (*W* = 26, *P* = 0.92) of PTP were unaffected by A/C in SynC-transduced neurons. Scale bars, 50 pA, 100 ms.

**Extended Data Fig. 3.**
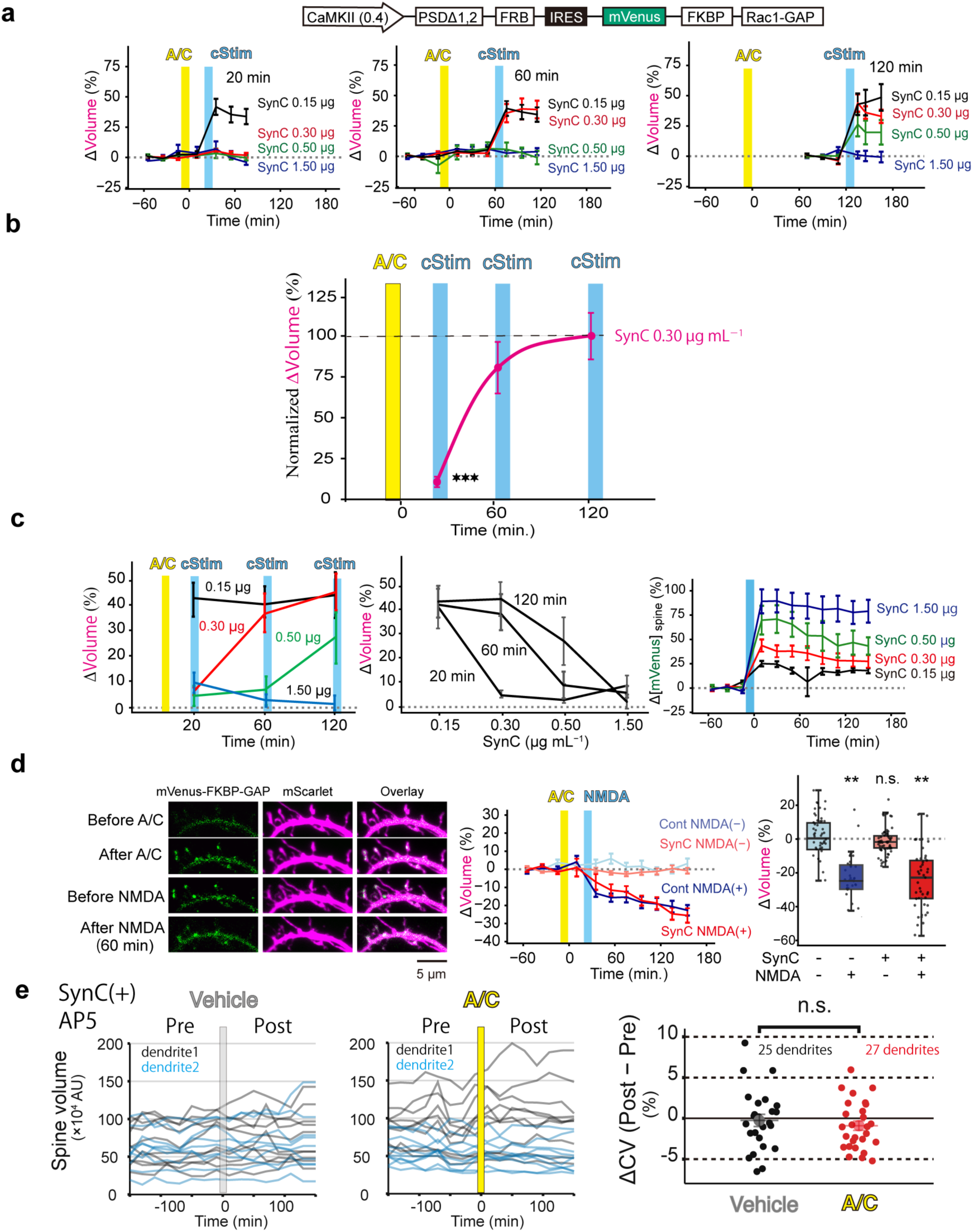
Functional specificity and reversibility of bicistronic SynC. **a**, Bicistronic SynC construct (top) and its action on average spine enlargement at cStim applied 20 min, 60 min, and 120 min after A/C treatment in cells lipofected with 1.5, 0.5, 0.3, or 0.15 µg mL^−1^ plasmids. **b**, Peak amplitudes of spine enlargement at three time points after A/C. Data were obtained for 0.3 µg mL^−1^ SynC plasmids from **c**. Mann-Whitney U test, *U* = 8, *P* = 2.81 × 10^−6^ at 20 min after A/C. Amplitudes were normalized to 100% (dashed line) relative to those without SynC at 120 min after A/C (Extended Data Fig. 1g). **c**, Time- and concentration-dependent blockade of spine enlargement (left, middle) and translocation (right) of SynC-mVenus following transduction with 1.5, 0.5, 0.3, or 0.15 µg mL^−1^ SynC-fusion plasmids. For 1.5 µg mL^−1^, 20, 60, 120 dendrites, 135, 144, 120 spines, 3, 4, 3 cultures (20, 60, 120 min). For 0.5 µg mL^−1^, 20, 60, 120 dendrites, 162, 145, 111 spines, 3, 4, 2 cultures. For 0.3 µg mL^−1^, 20, 60, 120 dendrites, 143, 148, 115 spines, 3, 4, 3 cultures. For 0.15 µg mL^−1^, 17, 17, 12 dendrites, 141, 109, 89 spines, 3, 3, 3 cultures. Relative translocation depended on SynC concentration because PSDΔ1,2-FRB levels varied accordingly (right). **d**, (Left) Example images of spine shrinkage induced by NMDA (20 µM NMDA) after A/C treatment in neurons lipofected with or without SynC-fusion (0.5 µg mL^−1^). Muscimol (400 nM) was continuously superfused to promote shrinkage. (Middle) Time courses of spine-volume changes induced by the NMDA protocol in the presence of A/C. *n* = 20 dendrites (206 spines, 6 cells, 3 cultures) for non-transfected cells without NMDA; *n* = 20 dendrites (157 spines, 8 cells, 4 cultures) for non-transfected cells with NMDA; *n* = 29 dendrites (270 spines, 9 cells, 3 cultures) for SynC-fusion-transfected cells without NMDA; *n* = 22 dendrites (194 spines, 8 cells, 5 cultures) with NMDA. (Right) Kruskal–Wallis test (*H*(3) = 43.92, *P* = 1.57 × 10^−9^) followed by post-hoc Steel test: *P* = 2.00 × 10^−5^ for non-transfected cells with NMDA; *P* = 0.365 for transfected cells without NMDA; *P* = 2.50 × 10^−5^ for transfected cells with NMDA. Scale bar, 5 µm. **e**, Spine-volume fluctuations in SynC-expressing cultures in the presence of AP5 (50 µM) treated with vehicle or A/C, from dendrites different from those in Fig. 1g. This condition controls for time-dependent drift during imaging. Left, middle, representative traces from two dendrites from vehicle (25 dendrites, 248 spines, 9 cells, 4 cultures) and A/C (27 dendrites, 270 spines, 10 cells, 4 cultures). Right, the change in CV (ΔCV, before vs after) compared between vehicle- and A/C-treated cultures. Welch’s *t*-test, *t* = −0.68, *P* = 0.50.

**Extended Data Fig. 4.**
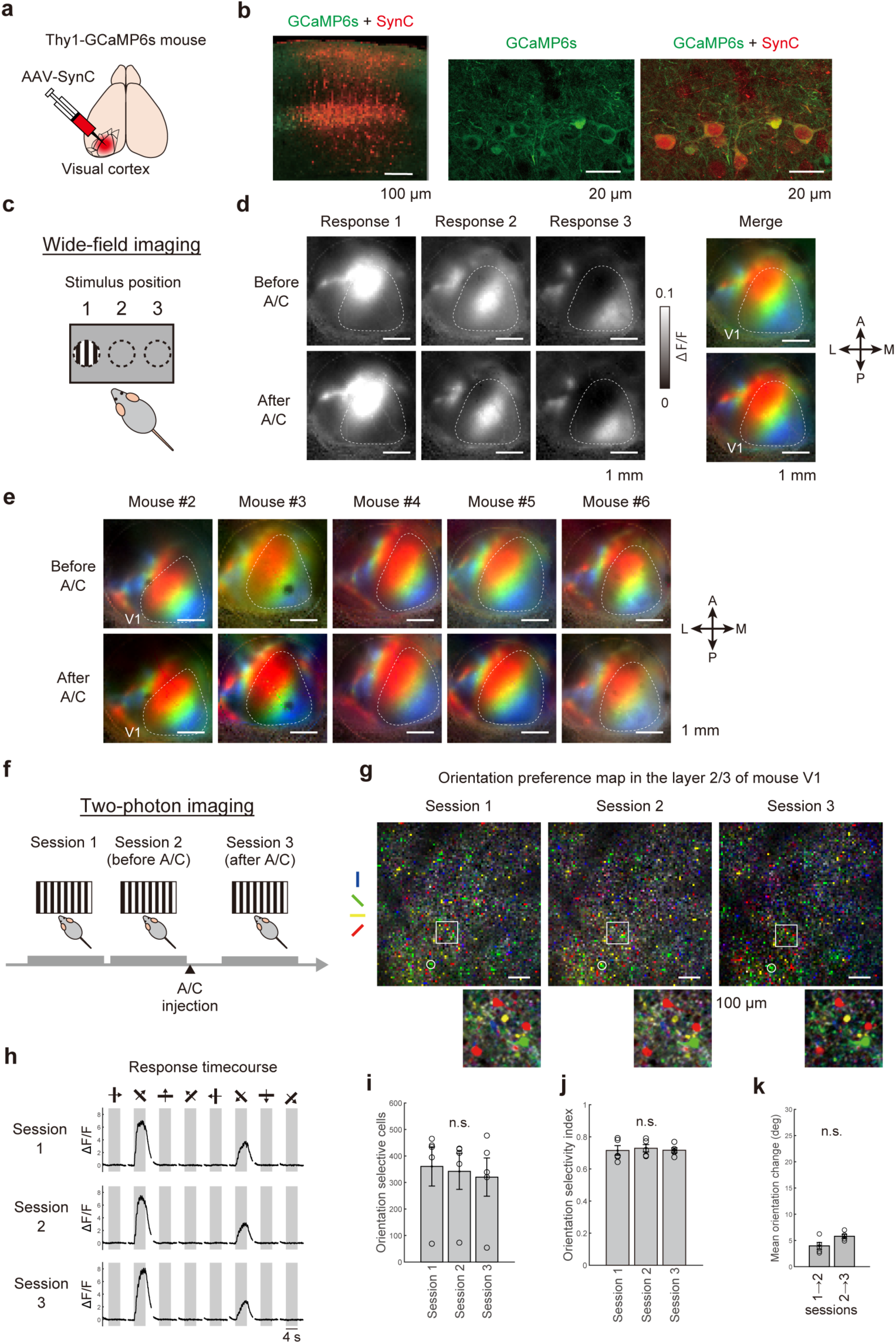
SynC expression and functional stability in mouse primary visual cortex (V1). **a**, Schematic of AAV::SynC injection into V1 of Thy1-GCaMP6s mice. **b**, Histological verification of GCaMP6s (green) and SynC (red) expression in V1. Scale bars, 100 µm, 20 µm, 20 µm. **c**, Wide-field calcium imaging for retinotopic mapping. Drifting grating stimuli (40° diameter) were presented at three horizontal positions. **d**, Representative response maps (ΔF/F) to the three stimuli (left) and merged retinotopic maps (right) before (top) and after (bottom) i.p. A/C. Scale bars, 1 mm. **e**, Retinotopic organization across six mice before and after i.p. A/C, total 6 mice with **h**. Scale bar, 1 µm. **f**, Timeline for two-photon calcium imaging in V1 across three sessions to evaluate tuning stability and the effect of i.p. A/C. **g**, Orientation-preference maps in layer 2/3 of mouse V1 for sessions 1–3 (top) and magnified views of the indicated regions (bottom). Each cell is coloured according to its preferred orientation, indicated by the four bars (blue, green, yellow and red). The circled cell is shown in **h**. Scale bars, 100 µm. **h**, Trial-averaged calcium traces from a representative neuron (circled in **g**). Grey shading indicates stimulus presentation (4 s). **i**, Number of orientation-selective cells across sessions (*n* = 5 mice; Friedman test, χ^2^(2) = 3.60, *P* = 0.165). **j**, Mean orientation selectivity index (OSI) across sessions (*n* = 5 mice; Friedman test, χ^2^(2) = 0.40, *P* = 0.819). **k**, Change in preferred orientation across sessions. The change in preferred orientation did not differ significantly between the control interval (sessions 1–2) and the post-A/C interval (sessions 2–3) (*n* = 5 mice; Wilcoxon signed-rank test, *P* = 0.125). Bars indicate mean ± s.e.m.

**Extended Data Fig. 5.**
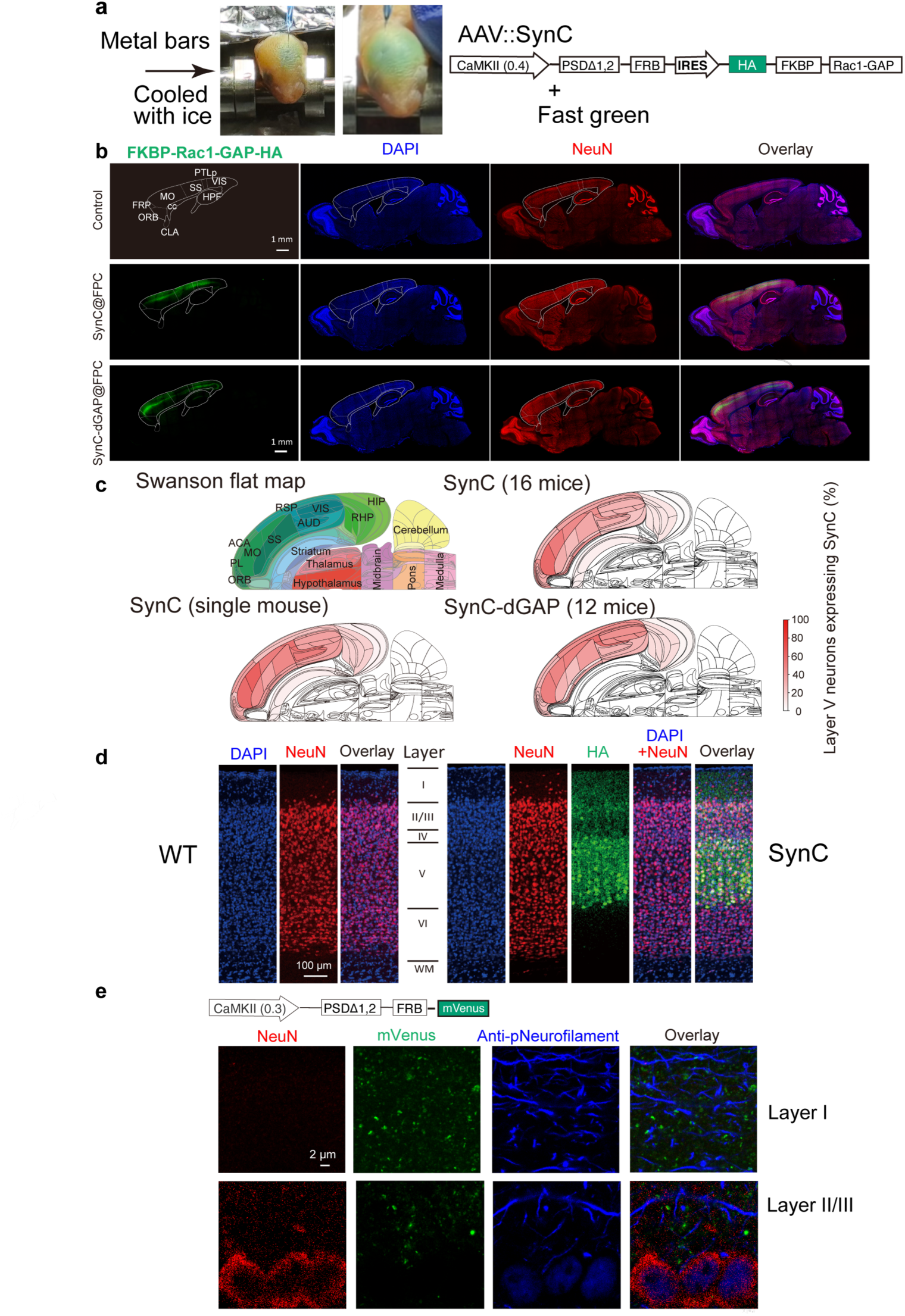
Anatomical distribution and cell-type specificity of SynC expression in the neocortex. **a**, Photographs of a neonatal pup immobilized between ice-cooled metal bars for hypothermic anaesthesia (left) and bicistronic AAV vector design (right). Under the CaMKII(0.4) promoter, the vector drives expression of PSDΔ1,2-FRB and HA-tagged FKBP-Rac1-GAP separated by an IRES. Fast green dye was used to visualize the injection site. **b**, Representative sagittal sections (1.35 mm lateral to midline) from non-injected control, SynC@FPC, and SynC-dGAP@FPC mice. Sections were immunolabelled for HA (green), NeuN (red), and nuclei (DAPI, blue). SynC signal was largely restricted to neocortex, whereas subcortical regions containing long-range cortical axons showed little HA immunoreactivity, consistent with limited axonal transport of the GAP module. Scale bars, 1 mm. **c**, Quantitative mapping of SynC expression across neocortex. Swanson flat map^61^ indicating cortical regions (top left). Anatomical abbreviations in **b**,**c** follow the Allen Mouse Brain Common Coordinate Framework^58^: ACA, anterior cingulate area; AUD, auditory areas; cc, corpus callosum; CLA, claustrum; FRP, frontal pole; HIP, hippocampal region; HPF, hippocampal formation; MO, somatomotor areas; ORB, orbital area; PL, prelimbic area; PTLp, posterior parietal association areas; RHP, retrohippocampal region; RSP, retrosplenial area; SS, somatosensory areas; VIS, visual areas. Pseudocolour heat maps showing the percentage of HA-positive layer V pyramidal neurons in the SynC group (top right, *n* = 16 mice), a single SynC mouse (bottom left), and the SynC-dGAP group (bottom right, *n* = 12 mice). **d**, Enlarged views of M1 lamination in WT and SynC@FPC mice across layers I–VI and white matter (WM). Immunolabelling of nuclei (DAPI, blue), NeuN (red) and their overlay for WT, and additionally, HA (green) for SynC@FPC. Scale bar, 100 µm. **e**, High-magnification images of layer I (top) and layer II/III (bottom) in mice expressing PSDΔ1,2-FRB-mVenus from a separate CaMKII(0.3) construct (top). NeuN (red), mVenus (green) and phospho-neurofilament (blue; axons). In layer I, mVenus-labelled spines lie adjacent to axons without overlapping axonal labelling; in layer II/III, mVenus is excluded from NeuN-positive somata and is concentrated in dendritic spines, consistent with postsynaptic restriction of the SynC anchor module. Fluorescence images are displayed with linear intensity scaling throughout. Scale bar, 2 µm.

**Extended Data Fig. 6.**
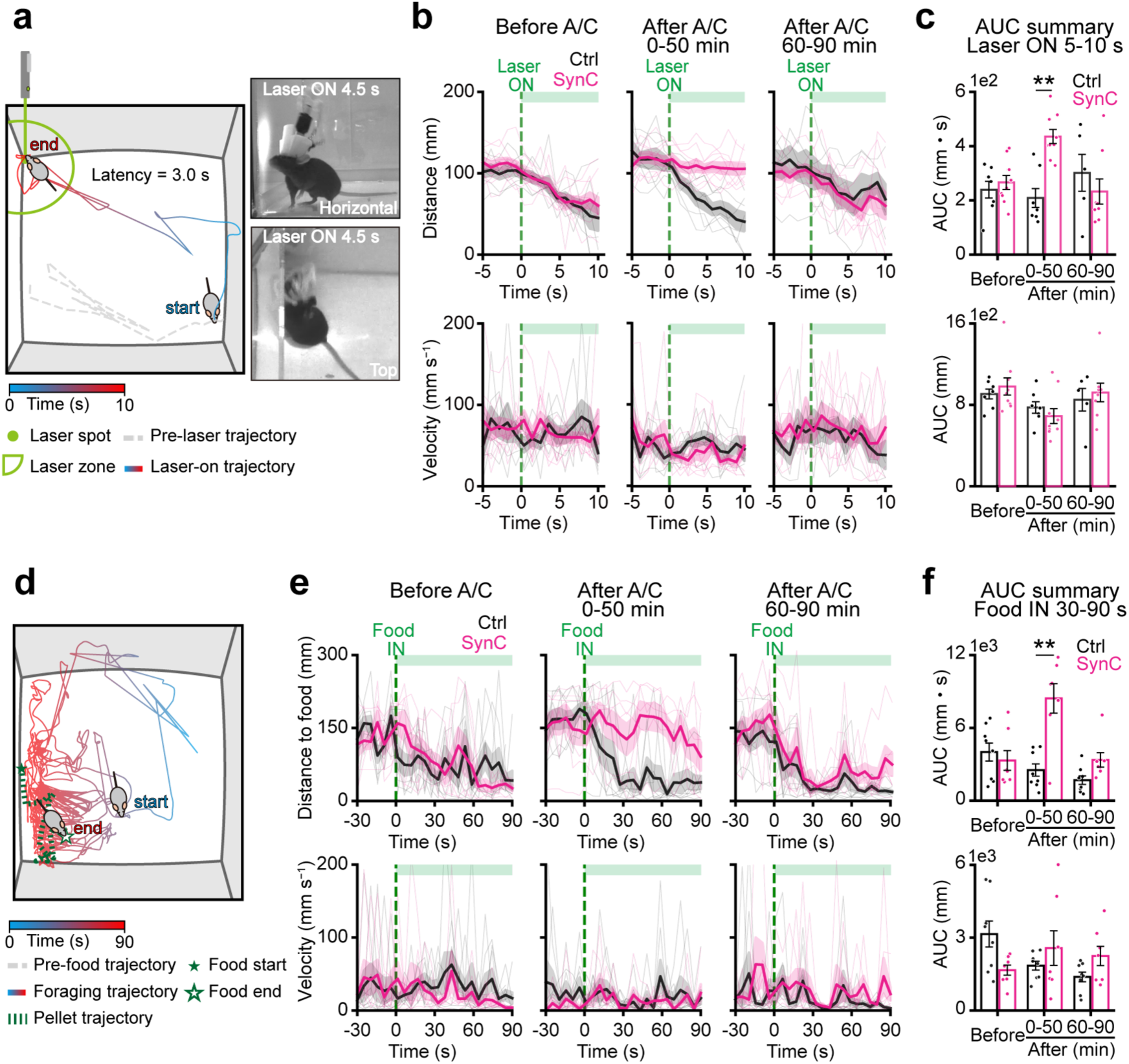
Quantitative analysis of laser-dot and food-pellet tasks. Quantification was performed during the dark (active) phase. **a**, Schematic illustration and image of experimental setup for the laser-dot task. The time after laser irradiation was pseudocolour-coded as indicated. **b**, The distance to the laser-corner (upper panels) and velocity of locomotion (lower panels) of wild-type (black) and SynC@FPC (magenta) mice before (*n* = 7 and 9), 0–50 min after (*n* = 7 and 9) and 60–90 min after (*n* = 5 and 8) i.p. A/C, respectively. **c**, Statistics of the data in b. Wild-type and SynC@FPC mice were compared with the Mann–Whitney U test on the AUC over 5–10 s after laser onset. The comparison at 0– 50 min after A/C was prespecified as the primary comparison; the remaining comparisons were secondary and are reported without multiplicity adjustment. Distance (upper panel): *U* = 28, *P* = 0.76 before i.p. A/C; *U* = 3, *P* = 0.0012, 0–50 min after (primary); *U* = 26, *P* = 0.44, 60–90 min after. Velocity (lower panel): *U* = 29, *P* = 0.84 before i.p. A/C; *U* = 42, *P* = 0.30, 0–50 min after; *U* = 20, *P* = 1.0, 60–90 min after. **d**, Schematic illustration and image of experimental setup for the food-pellet task. The time after laser irradiation was colour coded as indicated. The original (filled green star) and last (open green star) positions of the food pellet. Mice that entered State-C, as seen in the motion velocity in e, were excluded. **e**, The distance between the mouse and the food pellet (upper panels) and velocity of locomotion (lower panels) of wild-type (black) and SynC@FPC (magenta) mice before (*n* = 8 and 7), 0–50 min after (*n* = 8 and 7) and 60–90 min after (*n* = 8 and 7) i.p. A/C, respectively. **f**, Statistics of the data in e. Wild-type and SynC@FPC mice were compared with the Mann–Whitney U test on the AUC over 30–90 s after pellet introduction. The comparison at 0–50 min after A/C was prespecified as the primary comparison; the remaining comparisons were secondary and are reported without multiplicity adjustment. Distance (upper panel): *U* = 31, *P* = 0.78 before i.p. A/C; *U* = 6, *P* = 0.0093, 0–50 min after (primary); *U* = 9, *P* = 0.029, 60–90 min after. Velocity (lower panel): *U* = 43, *P* = 0.094 before i.p. A/C; *U* = 27, *P* = 0.96, 0–50 min after; *U* = 16, *P* = 0.19, 60–90 min after.

**Extended Data Fig. 7.**
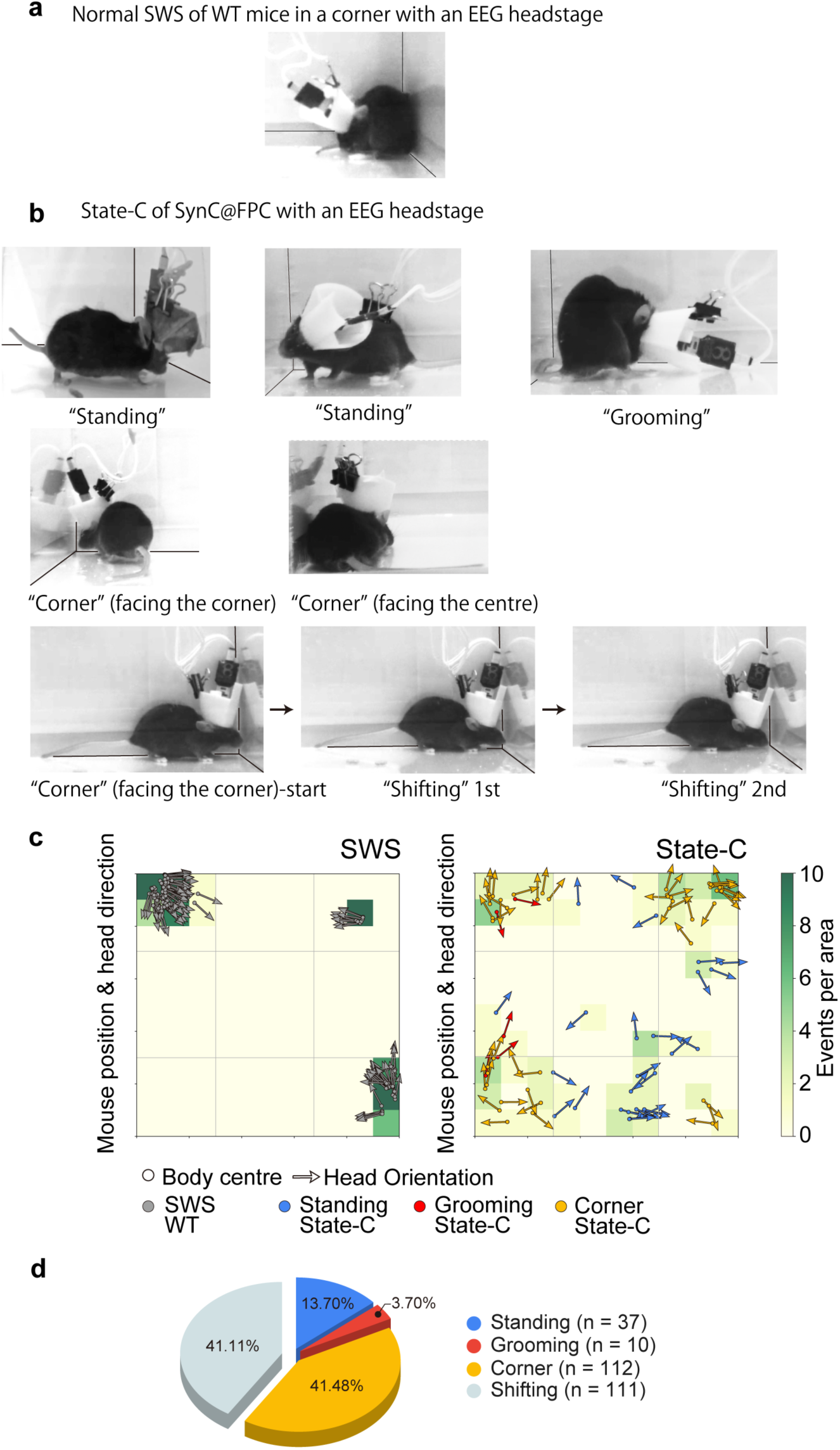
Behavioural postures and spatial distribution of State-C episodes. **a**, Representative image of a wild-type (WT) mouse with an EEG headstage during slow-wave sleep (SWS). Panels **a**–**c** are from a small open field (19.4 × 19.4 cm), whose close-up view allows postures to be classified offline from the recordings. **b**, Representative images of SynC@FPC mice with an EEG headstage during State-C illustrating four commonly observed postures: standing, grooming, corner, and shifting. Two examples are shown for standing, corner postures are shown facing either the corner or the centre, and shifting is shown as a three-frame sequence starting from a corner posture. In contrast to SWS, State-C episodes occurred during ongoing behaviour (e.g., locomotion, grooming, or corner navigation) and were identified by EEG-based state classification. **c**, Spatial distribution and head orientation during SWS (left; *n* = 6 mice, 112 epochs) and State-C (right; *n* = 5 mice, 80 epochs). Events per area are pseudocolour-coded; arrows indicate body-centre position and head orientation, colour-coded by posture (grey, SWS in WT; blue, standing; red, grooming; yellow, corner). Shifting episodes, which occur without a change in position, are not plotted. SWS episodes were concentrated near corners, whereas State-C episodes occurred across diverse locations and orientations, consistent with an abrupt behavioural arrest independent of position. **d**, Composition of State-C postures across identified epochs (*n* = 37 standing, 10 grooming, 112 corner and 111 shifting). Postures were identified by direct visual inspection in the large open field (35 × 35 cm).

**Extended Data Fig. 8.**
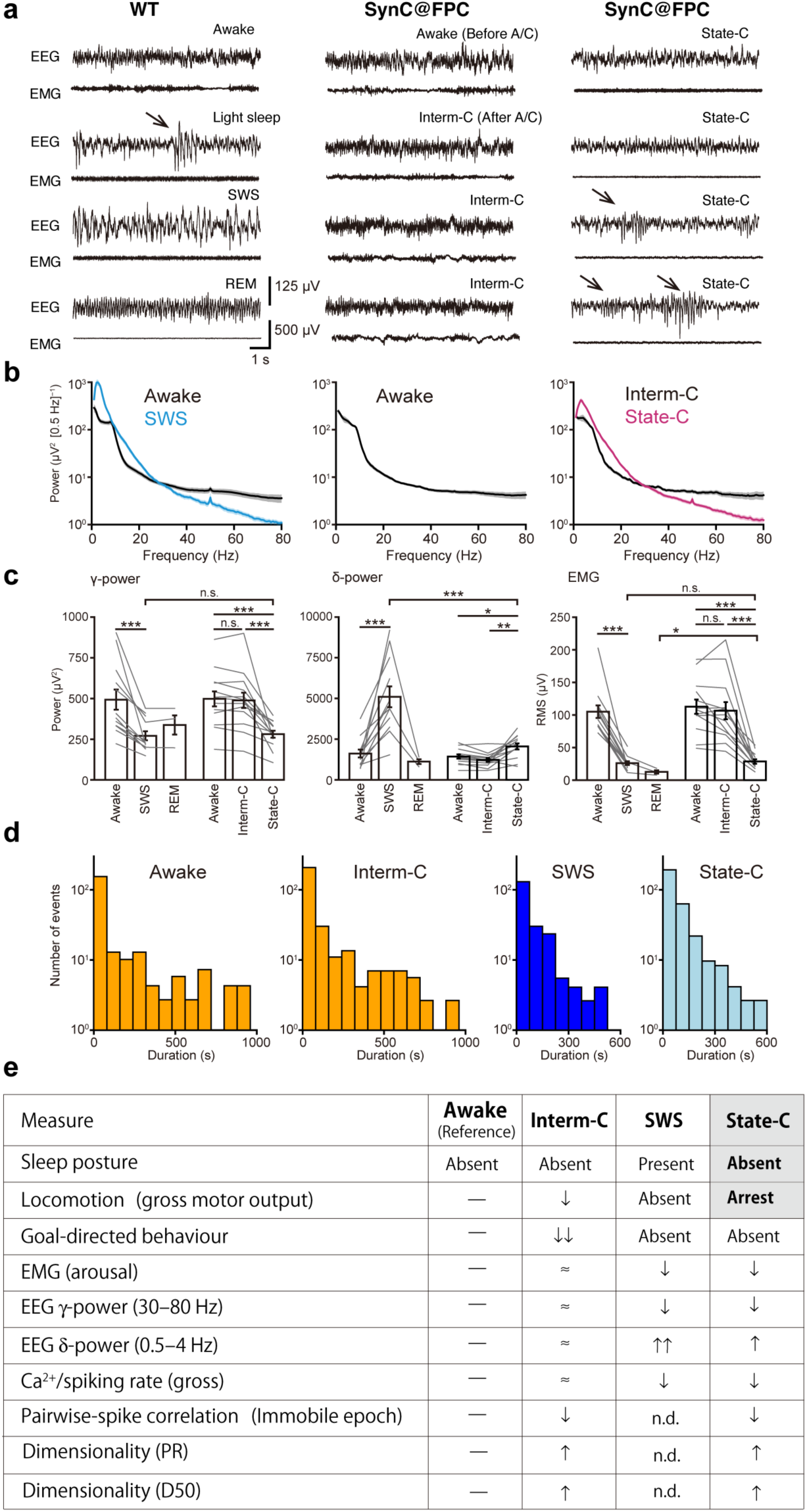
Comparative EEG and EMG features of awake, sleep, and SynC-induced states. **a**, Representative EEG and EMG traces. WT mice (left) show awake, light sleep, slow-wave sleep (SWS), and REM sleep. SynC@FPC mice (middle) show awake before A/C and Interm-C after A/C. Right, representative State-C episodes after A/C. Arrows indicate spindle-like events observed in both light sleep and State-C. Scale bars, 125 µV (EEG), 500 µV (EMG), and 1 s. **b**, EEG power spectra. Left, SWS (blue) vs awake (black) in WT mice. Middle, awake spectra in SynC@FPC mice before A/C. Right, State-C (magenta) vs Interm-C (black) after A/C. State-C shows reduced γ-band power without the sustained δ-band (0.5–4 Hz) elevation characteristic of SWS. **c**, Quantification of band power and muscle tone across states (*n* = 12 WT mice; *n* = 13 SynC@FPC mice). Within-genotype comparisons (Wilcoxon) were prespecified as primary and Bonferroni-corrected; between-genotype comparisons (Mann–Whitney) were secondary and are reported unadjusted. (Left) γ power (30–80 Hz) in Awake, SWS and REM (*n* = 12 WT mice; REM was observed in 3 of these mice) and in Awake, Interm-C and State-C (*n* = 13 SynC@FPC mice). Wilcoxon signed-rank test (WT, Awake vs SWS): *W* = 0, *P* = 4.9 × 10^−4^. For SynC@FPC, Friedman test (χ^2^(2) = 19.8, *P* = 4.9 × 10^−5^) followed by pairwise Wilcoxon signed-rank tests with Bonferroni correction: *W* = 28*, P* = 0.73 for Awake vs Interm-C; *W* = 0*, P* = 7.3 × 10^−4^ for Awake vs State-C; *W* = 0*, P* = 7.3 × 10^−4^ for Interm-C vs State-C. Mann-Whitney U test for State-C vs SWS: *U* = 66*, P* = 0.54. (Middle) δ-power (0.5–4 Hz) in Awake, SWS and REM (WT), and in Awake, Interm-C and State-C (SynC@FPC). Wilcoxon signed-rank tests (Awake vs SWS): *W* = 0, *P* = 4.9 × 10^−4^. For SynC@FPC, Friedman test (χ^2^(2) = 15.4, *P* = 4.5 × 10^−4)^ followed by pairwise Wilcoxon signed-rank tests with Bonferroni correction: *W* = 7*, P* = 1.4 × 10^−2^ for Awake vs State-C; *W* = 4*, P* = 5.1 × 10^−3^ *for Interm-C vs State-C*. Mann–Whitney U test for SWS vs State-C: *U* = 145*, P* = 7.5 × 10^−5^. (Right) EMG RMS amplitudes in Awake, SWS and REM (WT), and in Awake, Interm-C and State-C (SynC@FPC). Wilcoxon signed-rank tests (Awake vs SWS): *W* = 0, *P* = 4.9 × 10^−4^. For SynC@FPC mice, Friedman test (χ^2^(2) = 19.8, *P* = 4.9 × 10^−5^) followed by pairwise Wilcoxon signed-rank tests with Bonferroni correction: *W* = 29*, P* = 0.82 for Awake vs Interm-C*; W* = 0*, P* = 7.3 × 10^−4^ for Interm-C vs State-C; *W* = 0*, P* = 7.3 × 10^−4^ for Awake vs State-C. Mann-Whitney U test: *U* = 78, *P* = 1.0 for SWS vs State-C; *U* = 2*, P* = 0.0143 for State-C vs REM (*n* = 13 and 3). Bars show mean ± s.e.m.; grey lines, individual mice. **d**, Episode-duration distributions for awake (*n* = 12 mice), Interm-C (*n* = 13), SWS (*n* = 12), and State-C (*n* = 13). Histograms show the number of episodes per duration bin; note the logarithmic ordinate. **e**, State-dependent behavioural and neural signatures (normalized to Awake). Symbols denote changes relative to Awake (reference): ≈ no detectable change; ↑/↓ increase/decrease; ↑↑/↓↓ marked change; n.d., not determined: local somatic Ca^2+^ imaging of pyramidal neurons does not provide a direct readout of the electrophysiological synchrony that defines SWS^38^, and sustained SWS was too rare during treadmill imaging for reliable correlation and dimensionality estimates. **Summary of state definitions: State-C, abrupt** behavioural arrest after A/C lasting ≥10 s without sleep posture; accompanied by γ ↓, EMG ↓, δ slightly ↑, mean activity ↓, pairwise correlation ↓ and dimensionality (D50, PR) ↑. **Interm-C, post-A/C** periods outside State-C, before recovery; accompanied by γ ≈, EMG ≈ and mean activity ≈, with pairwise correlation ↓, dimensionality (D50, PR) ↑ and goal-directed behaviour ↓↓. D50, number of principal components explaining 50% of variance; PR, participation ratio. **Data sources:** behavioural tasks (Fig. 3,4g, Extended Data Fig. 6), EEG/EMG (Fig. 4; Extended Data Fig. 8a–d), Ca^2+^/spiking and correlations (Fig. 5), dimensionality (Extended Data Fig. 9).

**Extended Data Fig. 9.**
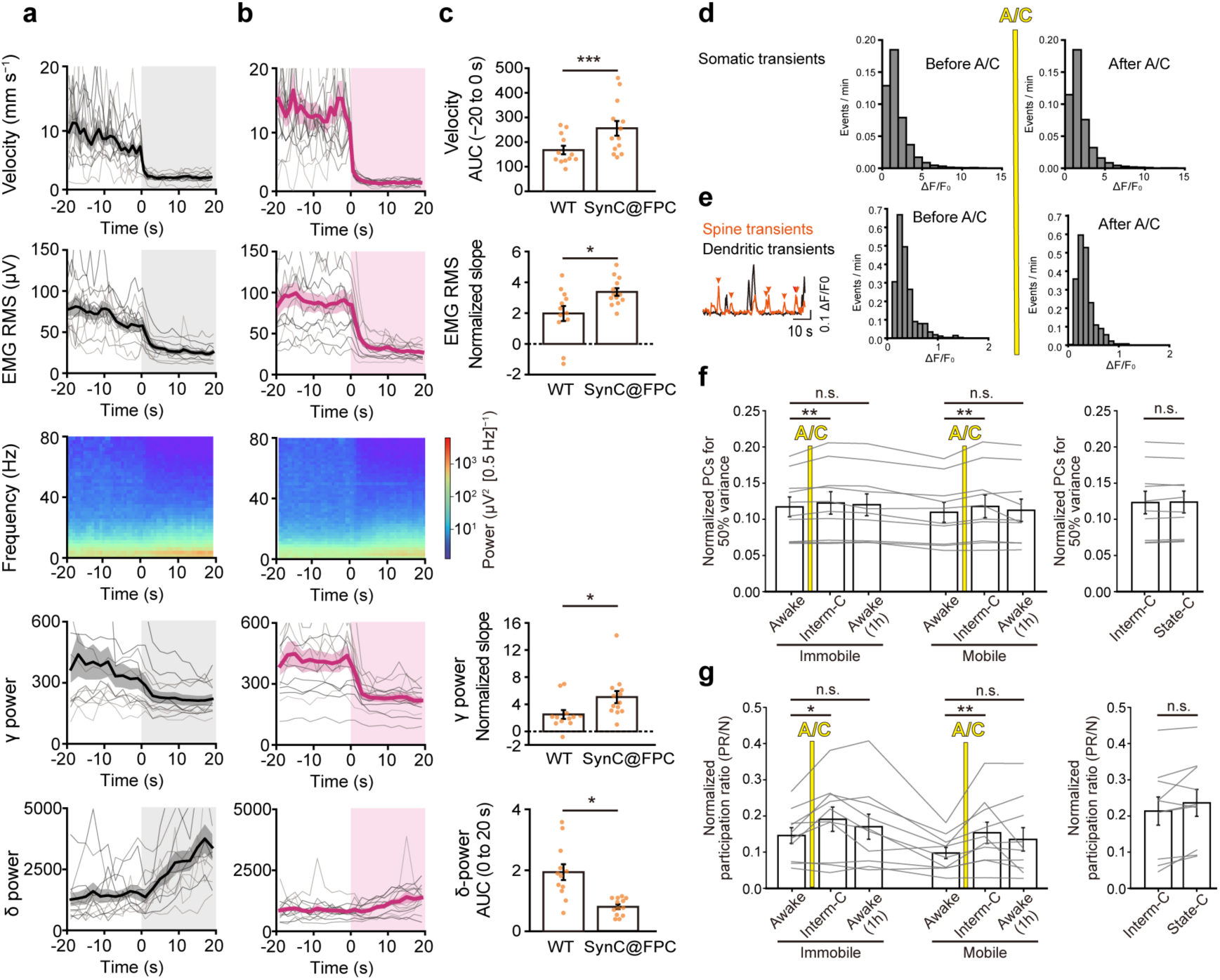
Transitions into State-C, stability of Ca^2+^ signals, and network dimensionality. Recordings were performed during the dark (active) phase. **a**,**b**, Event-aligned dynamics of behavioural and physiological parameters (top to bottom: locomotor velocity, EMG RMS, EEG spectrogram, and γ-band [30–80 Hz] and δ-band [0.5–4 Hz] power). Transitions were aligned to onset (*t* = 0) of Awake→SWS in WT mice (**a**; *n* = 12) and Interm-C→State-C in SynC@FPC mice (**b**; *n* = 13) within 2 h after i.p. A/C. Thin grey traces show per-mouse means (3–16 events per mouse); thick coloured traces show across-mouse means. **c**, Quantification of transition dynamics, arranged in the same row order as a,b. Two measures were used: the area under the curve (AUC), and a normalized slope as an index of abruptness (slope over −3 to +1 s divided by slope over −10 to +10 s). WT and SynC@FPC mice were compared by Mann–Whitney tests: velocity AUC (−20 to 0 s), *U* = 143*, P* = 1.4 × 10^−4^; EMG RMS normalized slope, *U* = 35, *P* = 0.019; γ-band normalized slope, *U* = 32, *P* = 0.011; δ-band AUC (0 to 20 s), *U* = 32, *P* = 0.011. The γ-band comparison was prespecified as the primary comparison and the remaining comparisons as secondary, the latter reported without multiplicity adjustment. Error bars, s.e.m. **d**, Distributions of detected somatic GCaMP7f event amplitudes (ΔF/F_0_) per minute before (left) and after (right) A/C heterodimerizer administration. *n* = 9 SynC@FPC mice (5,136 and 6,895 events). Mean amplitudes were 1.94 ± 1.06 and 1.98 ± 1.70 (mean ± s.d.) before and after A/C — a difference of 0.03 s.d., with the two distributions essentially superimposed. **e**, Left, spine Ca**^2+^** transients (red trace) isolated from the signal of the adjacent dendritic shaft (black trace); red arrowheads mark detected spine transients. Scale bars, 0.1 ΔF/F_0_ and 10 s. Right, distributions of detected spine-transient amplitudes (ΔF/F_0_) per minute before and after A/C. *n* = 52 spines from 3 SynC@FPC mice (550 events before and 537 events after). Mean amplitudes were 0.374 ± 0.24 and 0.352 ± 0.21 (mean ± s.d.) before and after A/C — a difference of 0.10 s.d., with the two distributions essentially superimposed. **f**,**g**, Normalized number of principal components accounting for 50% of the total variance (D50/N, f) and normalized participation ratio (PR/N, **g**), derived from Ca^2+^ imaging in the same 10 SynC@FPC mice as in Fig. 5e,g and for the same phases. Grey lines, individual mice. Exact permutation test (immobile; mobile in parentheses) — D50: Interm-C vs Awake (100 s), *P* = 0.0078 (0.0020); Awake (1 h) vs Awake (100 s), *P* = 0.26 (0.10); immobile Interm-C vs State-C (50 s), *P* = 0.057. PR/N: Interm-C vs Awake, *P* = 0.025 (0.0020); Awake (1 h) vs Awake, *P* = 0.25 (0.13); immobile Interm-C vs State-C, *P* = 0.057. Fixed-sequence testing as in Fig. 5e.

**Extended Data Fig. 10.**
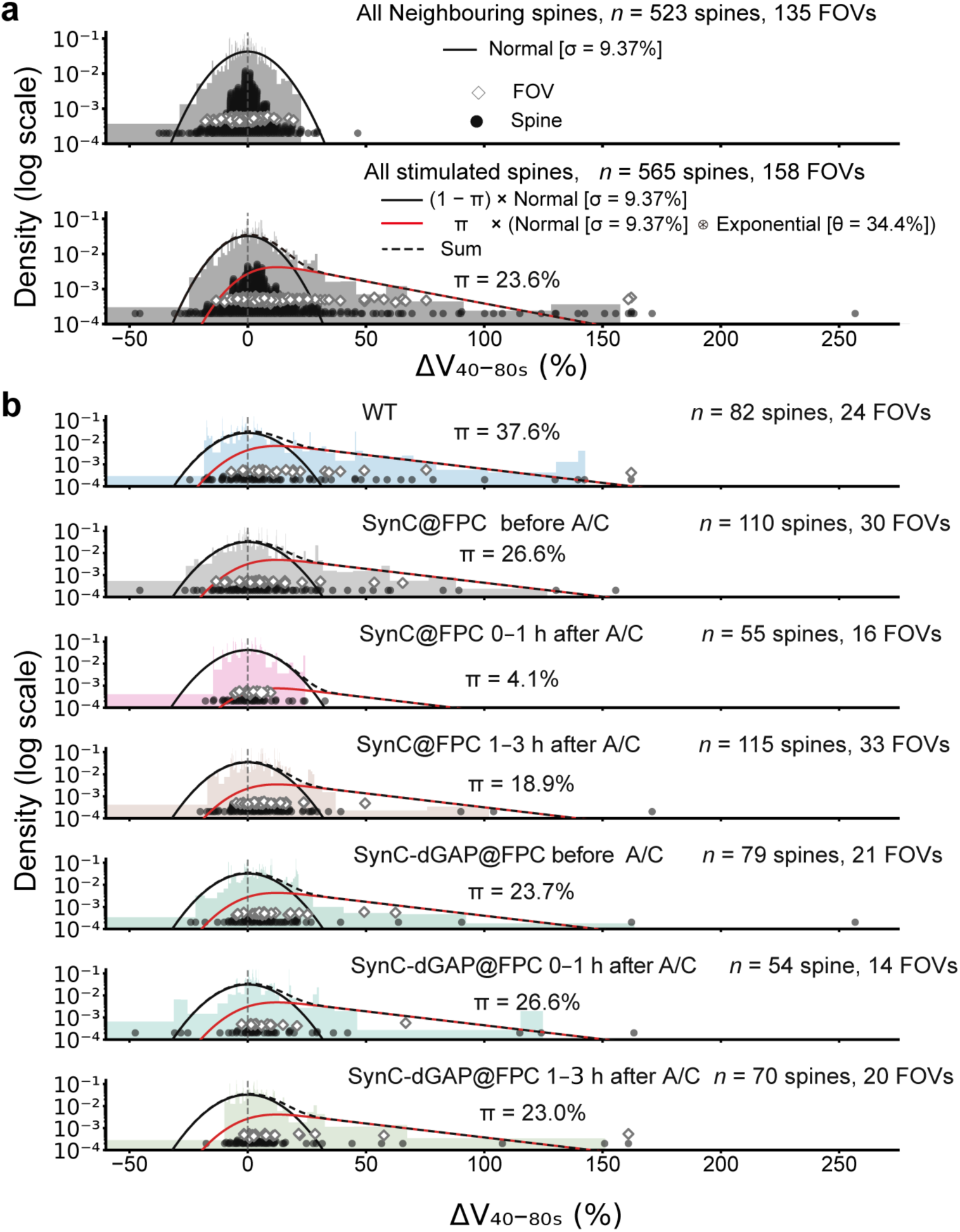
Distributional basis and condition dependence of the permissive spine fraction. **a,** Empirical probability-density distributions of mean ΔV over 40–80 s after stimulation for all stimulated spines (*n* = 565, bottom) and all neighbouring spines (*n* = 523, top), shown on logarithmic ordinates. Distributions are displayed as 1-percent percentograms: variable-width bins were defined independently for each distribution using approximately 1-percentile intervals, and bin heights were divided by bin width such that the total histogram area equalled one. For datasets containing fewer than 100 observations, the number of bins was limited to the number of observations so that every bin contained at least one observation. Black points and open diamonds indicate individual spine values and FOV means, respectively. The pooled neighbouring-spine data (top; 523 spines, 135 fields of view [FOVs], 9 mice) defined a zero-centred normal distribution with a maximum-likelihood σ of 9.37%. The pooled stimulated-spine data (bottom; 565 spines, 158 FOVs, 11 mice) were fitted with a two-component mixture in which a fraction π of spines shows an exponentially distributed enlargement (θ = 34.4%) observed with the same normal measurement noise (σ = 9.37%), and the remaining (1 − π) show noise alone; the fitted permissive fraction was π = 23.6% (pooled across spines). Dashed line, sum of the two components. **b,** Condition-specific variable-width histograms of stimulated-spine mean ΔV over 40– 80 s, shown on logarithmic probability-density ordinates as in **a**. Data are shown for WT, SynC before A/C, SynC 0–1 h after A/C, SynC 1–3 h after A/C, SynC-dGAP before A/C, SynC-dGAP 0–1 h after A/C and SynC-dGAP 1–3 h after A/C (cohorts and n as in Fig. 6d–g). The pooled normal and exponential distribution were shared across conditions, whereas the fitted permissive fraction, π, was condition-specific. Estimated π values were 37.6%, 26.6%, 4.1%, 18.9%, 23.7%, 26.6% and 23.0%, respectively. The permissive component was required in all conditions except after A/C (π > 0; spine-level parametric bootstrap, *P* < 0.001), where it was much reduced (*P* = 0.013). Between-condition comparisons are shown in Fig. 6h. Black points indicate individual spines, and open diamonds indicate FOV means.

**Extended Data Movie 1. Laser-dot task (WT mouse)**

A wild-type (z233) mouse wearing an EEG headset chased a laser dot projected onto the corner of a small open field (19.4 × 19.4 cm) within 10 s. The laser was directed toward the corner to avoid reflections, although mice normally chase the laser dot even on a flat surface. The movie is shown at 2× real time.

**Extended Data Movie 2. Laser-dot task (SynC@FPC mouse before i.p. A/C)**

A SynC@FPC mouse (z244-5, 1117 s before A/C) expressing SynC broadly in the frontoparietal cortex chased the laser dot within 10 s, similarly to the WT mouse. The movie is shown at 2 × real time.

**Extended Data Movie 3. Laser-dot task (SynC@FPC mouse after i.p. A/C)**

The same SynC@FPC mouse (z244-5, 2298 s after A/C) as in Movie 2. The mouse failed to chase the laser dot within 10 s. The movie is shown at 2× real time.

**Extended Data Movie 4. Laser-dot task (SynC@FPC mouse 1 h after i.p. A/C)**

The same SynC@FPC mouse (z244-5, 3483 s after A/C) as in Movie 2 and 3. The mouse again chased the laser dot within 10 s, indicating recovery of normal behaviour. The movie is shown at 2× real time.

**Extended Data Movie 5. Food-pellet task (WT mouse after i.p. A/C)**

A WT mouse (z239, 785 s after A/C) grasped and ate a food pellet placed in the arena within 30 s. The movie is shown at 2× real time.

**Extended Data Movie 6. Food-pellet task (SynC@FPC mouse after i.p. A/C)**

A SynC@FPC mouse (z227-6, 1,203 s after A/C) repeatedly passed over the food pellet without eating and eventually began consuming it only after approximately 5 min. The mouse spent an unusually long period grooming near the pellet and did not respond to the laser dot. Once it grasped the pellet, however, it ate normally, indicating that consummatory behaviour remained intact despite impaired feeding initiation. The movie is shown at 2× real time.

**Extended Data Movie 7. Food-pellet task (SynC@FPC mouse 1 h after i.p. A/C)**

The same mouse (z227-6, 5204 s after A/C) as in Movie 6. The mouse rapidly approached, grasped, and ate the food pellet within 10 s. The movie is shown at 2× real time.

**Extended Data Movie 8. “Standing” posture example 1 during State-C (SynC@FPC mouse)**

A SynC@FPC mouse (z156-3, 444 s after A/C) exhibited a “standing” posture upon behavioural arrest during running, shown from side and top views in a small (15 × 15 cm) open field. EEG, EMG, and velocity traces are displayed simultaneously. The movie is shown in real time.

**Extended Data Movie 9. “Standing” posture example 2 during State-C (SynC@FPC mouse)**

A SynC@FPC mouse (z248-2, 1007 s after A/C) exhibited a “standing” posture upon behavioural arrest during running, shown from side and top views in a small (19.4 × 19.4 cm) open field. EEG, EMG, and velocity traces are displayed simultaneously. The movie is shown in real time.

**Extended Data Movie 10. “Grooming” posture during State-C (SynC@FPC mouse)**

A SynC@FPC mouse (z248-1, 1873 s after A/C) exhibited a “grooming” posture upon behavioural arrest during grooming, shown from side and top views in a small open field. EEG, EMG, and velocity traces are displayed simultaneously. The movie is shown in real time.

**Extended Data Movie 11. “Corner” posture during State-C (SynC@FPC mouse)**

A SynC@FPC mouse (z248-2, 1241 s after A/C) exhibited a “corner” posture upon behavioural arrest while approaching a corner in the small open field. EEG, EMG, and velocity traces are displayed simultaneously. The movie is shown in real time.

**Extended Data Movie 12. “Shifting” posture during State-C (SynC@FPC mouse)**

A SynC@FPC mouse (z244-2) exhibited brief “shifting” movements at 115 s and 225 s following a corner posture, coinciding with short bouts (0.5 s and 2 s) of Interm-C. EEG, EMG, and velocity traces are displayed simultaneously. The movie is shown in real time.

Extended Data Movie 13. Normal sleep-preparation behaviour (WT mouse)

Before sleep onset, a WT mouse (z155-2, 1198 s after A/C) slowly walked toward a corner as EEG γ-power was already decreasing. The mouse then turned to face the centre, curling its body and tail. EEG δ-power increased after the mouse adopted a full sleep posture. The movie is shown in real time.

**Extended Data Movie 14. Transition from Interm-C to State-C (SynC@FPC mouse)**

A SynC@FPC mouse (z178-4, 1415 s after A/C) walked toward a corner and abruptly entered behavioural arrest precisely when EEG γ-power declined sharply. EEG δ-power showed only a slight increase. Spindle waves appeared shortly after the onset of State-C in this case. The movie is shown in real time.

**Extended Data Movie 15. Treadmill running before i.p. A/C (SynC@FPC mouse)**

A mirror beneath the treadmill captured normal running and transient stalling movements of the SynC@FPC mouse (z251-3, 1478 s before A/C), whose hindlimbs were widely spaced, providing stable treadmill control. The movie is shown in real time.

**Extended Data Movie 16. Treadmill running after i.p. A/C (SynC@FPC mouse)**

The same SynC@FPC mouse (z251-3, 614 s after A/C) as in Extended Data Movie 15 showed somewhat clumsier running (hindlimbs narrowly spaced) and occasional stalling, failing to adequately control the treadmill. The movie is shown in real time.

