## Supplementary Table 1 for "Rapid associative spine enlargement is required for cognitive function and stable wakefulness"

| State | Behaviour | EEG | EMG | Autonomic |
|---|---|---|---|---|
| Awake<br>(reference) | Exploration; goal-directed | γ√; δ low;<br>Low-amp fast | High | HR high; resp ≈;<br>pupil dilated |
| **Interm-C**<br>(this study) | Locomotor-active;<br>**goal-directed control ↓** | **γ√; δ low;**<br>low-amp fast | **High** | HR high; resp ≈;<br>pupil dilated |
| **State-C**<br>(this study) | **Abrupt behavioural arrest with eyes open and non-sleep posture,**<br>with exponentially distributed bouts, as in SWS | **γ↓↓; small δ↑; isolated spindles; no SWD** | **Low** | Mild HR↓; resp ≈ ;<br>miosis |
| Light sleep[1] | **Sleep posture**;<br>short in mouse | **γ↓↓; small δ↑;**<br>occasional spindles | **Low** | Mild HR↓; resp ≈ ;<br>miosis |
| SWS<br>(deep NREM)[1] | **Sleep posture** | **γ↓↓; δ↑↑;<br>high-amp;** | **Low** | Mild HR↓; resp ≈ ;<br>miosis |
| REM sleep[2] | Sleep posture | γ↓; **θ-dominant;**<br>low-amp fast | **Lowest** | Variable |
| Freezing<br>(defensive)[3] | **Immobile awake** | **γ√; δ low;** | Moderate–<br>high | HR ↓ |
| Chx10-PPN-induced motor arrest[4] | **Abrupt pause-and-play arrest** | n.d. | Ongoing pattern held | Mild HR↓; **resp↓↓ (apnea)** |
| Absence seizure[5] | Behavioural arrest;<br>brief bouts | γ↓↓; **SWD** | Low–<br>moderate | Variable |
| Cataplexy in narcolepsy[6] | Emotion-triggered collapse | γ↓↓; **REM-like (θ-dominant);**<br>rapid transitions | **Very low** | Variable |
| General anaesthesia (propofol)[7] | **Lying down** | γ↓↓; δ↑ ;<br>**frontal α–coherence** | Very low | Autonomic suppression |
| Catalepsy<br>(neuroleptic-induced)[8] | **Rigid immobility with imposed posture maintenance** | γ↓; often δ↑ | **High** | Not defining |

### Supplementary Table 1. Differential features distinguishing Interm-C and State-C arrest from canonical sleep states, seizure, anaesthesia and arrest-like states.

Entries labelled "this study" were directly observed/defined here. **Notes:** γ, 30–80 Hz; δ, 0.5–4 Hz; θ, 4–8 Hz; SWD, spike-and-wave discharges. See Methods for frequency band definitions and statistics. Upper section summarizes states observed in this study (including Interm-C/State-C and baselines); lower section lists canonical comparator states used for differential classification. Shading indicates the domains most characteristic of each state; bold indicates the defining features within those domains.
